# A global atlas of substrate specificities for the human serine/threonine kinome

**DOI:** 10.1101/2022.05.22.492882

**Authors:** Jared L. Johnson, Tomer M. Yaron, Emily M. Huntsman, Alexander Kerelsky, Junho Song, Amit Regev, Ting-Yu Lin, Katarina Liberatore, Daniel M. Cizin, Benjamin M. Cohen, Neil Vasan, Yilun Ma, Konstantin Krismer, Jaylissa Torres Robles, Bert van de Kooij, Anne E. van Vlimmeren, Nicole Andrée-Busch, Norbert Käufer, Maxim V. Dorovkov, Alexey G. Ryazanov, Yuichiro Takagi, Edward R. Kastenhuber, Marcus D. Goncalves, Olivier Elemento, Dylan J. Taatjes, Alexandre Maucuer, Akio Yamashita, Alexei Degterev, Rune Linding, John Blenis, Peter V. Hornbeck, Benjamin E. Turk, Michael B. Yaffe, Lewis C. Cantley

## Abstract

Protein phosphorylation is one of the most widespread post-translational modifications in biology. With the advent of mass spectrometry-based phosphoproteomics, more than 200,000 sites of serine and threonine phosphorylation have been reported, of which several thousand have been associated with human diseases and biological processes. For the vast majority of phosphorylation events, it is not yet known which of the more than 300 protein Ser/Thr kinases encoded in the human genome is responsible. Here, we utilize synthetic peptide libraries to profile the substrate sequence specificity of nearly every functional human Ser/Thr kinase. Viewed in its entirety, the substrate specificity of the kinome was substantially more diverse than expected and was driven extensively by negative selectivity. Our kinome-wide dataset was used to computationally annotate and identify the most likely protein kinases for every reported phosphorylation site in the human Ser/Thr phosphoproteome. For the small minority of phosphosites where the protein kinases involved have been previously identified, our predictions were in excellent agreement. When this approach was applied to examine the signaling response of tissues and cell lines to hormones, growth factors, targeted inhibitors, and environmental or genetic perturbations, it revealed unexpected insights into pathway complexity and compensation. Overall, these studies reveal the full extent of substrate specificity of the human Ser/Thr kinome, illuminate cellular signaling responses, and provide a rich resource to link unannotated phosphorylation events to biological pathways.

## INTRODUCTION

Phosphorylation of proteins on serine, threonine, tyrosine, and histidine residues controls nearly every aspect of eukaryotic cellular function^1–4^. Misregulation of protein kinase signaling commonly results in human disease^5–8^. Deciphering the cellular roles of any protein kinase requires elucidation of its downstream effector substrates. The majority of kinase-substrate relationships that have been published to date, however, involve a relatively small number of well-studied protein kinases, while few, if any, substrates have been identified for the majority of the ~300 human protein Ser/Thr kinases within the human kinome^9–11^. This lack of knowledge of kinase-substrate relationships limits the interpretation of large mass spectrometry-based phosphoproteomic datasets, which to date have collectively reported over 200,000 Ser and Thr phosphorylation sites on human proteins^12–15^. The specific kinases responsible for these phosphorylation events have been reported for <4% of these sites^15^, severely limiting the understanding of cellular phosphorylation networks.

Well-studied serine/threonine kinases are generally known to recognize specific amino acid residues at multiple positions surrounding the site of phosphorylation^16–21^. This short linear motif, which is characteristic of a given protein kinase, ensures fidelity in signaling pathways regulating phosphorylation at a given Ser or Thr residue. Knowledge of kinase recognition motifs can facilitate discovery of new substrates, for example by scanning phosphoproteomics data for matching sequences. However, to date, phosphorylation site sequence motifs are known for only a subset of the human protein Ser/Thr kinome. In some cases, kinase recognition motifs have been inferred by alignment of known cellular phosphorylation sites that have been experimentally identified over many years. This process is slow and laborious and limited to kinases with large numbers of established substrates. We have previously described combinatorial peptide library screening methods that allow for rapid determination of specificity for individual kinases based on phosphorylation of peptide substrates^22,23^. Here, we apply those methods to experimentally determine the optimal substrate specificity for nearly the entire human serine-threonine kinome, characterize the relationship between kinases based on their motifs, and computationally utilize this data to identify the most likely protein kinase to phosphorylate any site identified by mass spectrometry or other techniques. Finally, we show how this information can be applied to capture complex changes in signaling pathways in cells and tissues following genetic, pharmacological, metabolic, and environmental perturbations.

## RESULTS

### Phosphorylation site substrate specificity of the human serine-threonine kinome

Substrate recognition motifs across the human Ser-Thr kinome were determined by performing positional scanning peptide array (PSPA) analysis. We used a previously reported combinatorial peptide library that systematically substitutes each of 22 amino acids (20 natural amino acids plus phospho-Thr and phospho-Tyr) at nine positions surrounding a central phospho-acceptor position containing equivalent amounts of Ser and Thr (Fig. 1a)^23^. Using purified recombinant kinase preparations, we successfully obtained phosphorylation site motifs for 303 Ser/Thr kinases, covering every branch of the human Ser/Thr kinase family tree as well as a collection of atypical protein kinases (Fig. 1b, Fig. S1). The large majority of these kinases, including 86 understudied “dark” kinases, had not been previously profiled.

**Fig. 1.**
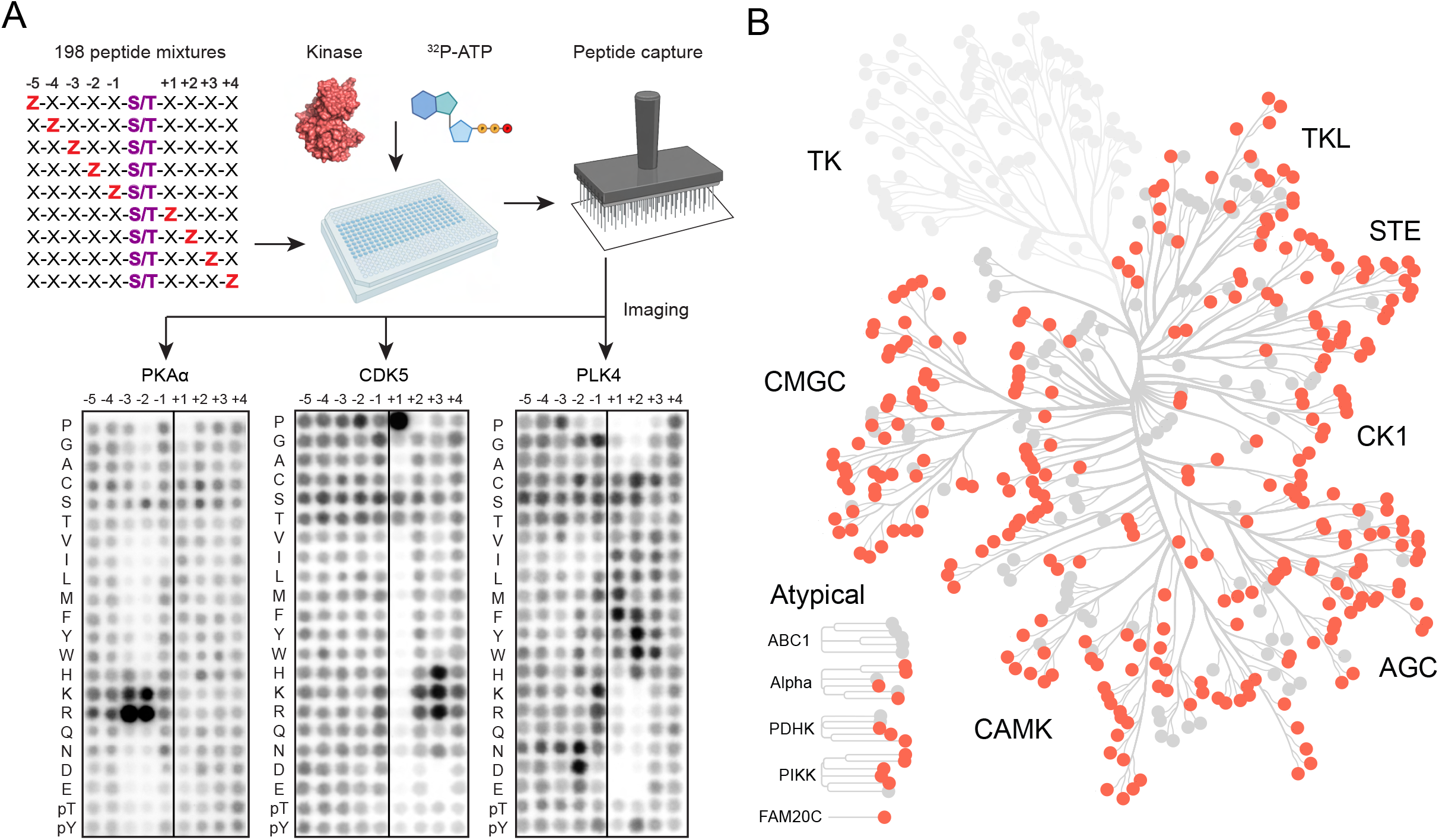
Profiling the substrate specificity of the human serine/threonine kinome. (A) Experimental workflow for positional scanning peptide arrays and representative results. (B) Dendrogram of the human protein kinome that highlights the serine/threonine kinases analyzed in this work.

Position-specific scoring matrices (PSSMs) derived from quantified PSPA data were analyzed by hierarchical clustering to compare kinase substrate motifs across the kinome (Fig. 2). As expected, kinases sharing substantial sequence identity displayed a high degree of similarity in their optimal substrate motifs. However, we found many cases where clustering by PSSM did not strictly recapitulate evolutionary phylogenetic relationships between kinases inferred from their primary sequences (Fig. 2). Instead, members of most major kinase groups were distributed throughout the dendrogram, reflecting numerous examples where kinases with low overall sequence identity have converged to phosphorylate similar optimal sequence motifs. For example, we found that a number of distantly related kinases (in the YANK, casein kinase 1 and 2, GRK, and TGF-β receptor families) converged to phosphorylate similar sequence motifs despite their very disparate locations on the kinome tree (Fig. 2, Cluster 3).

**Fig. 2.**
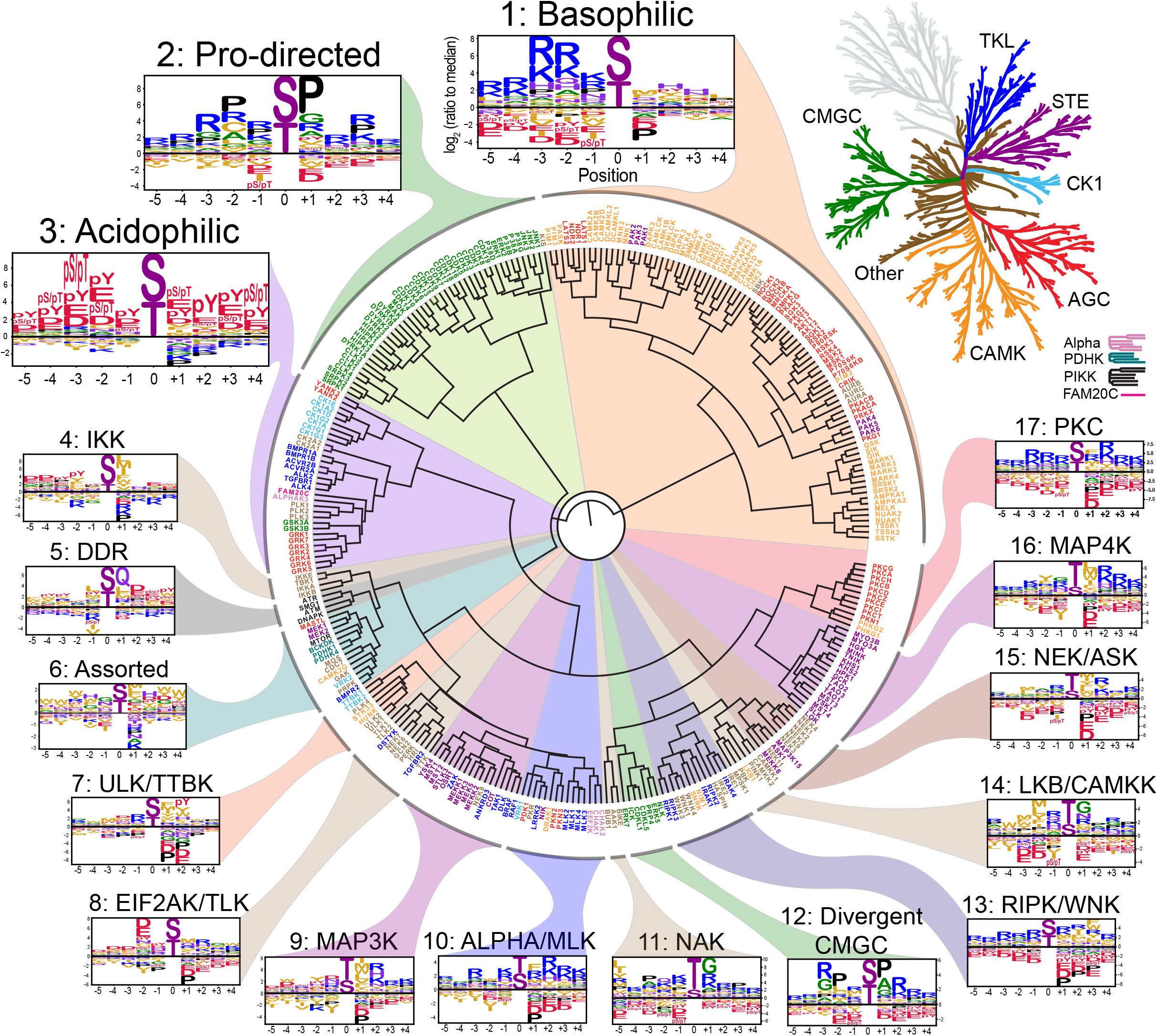
Phosphorylation site motif tree of the human Ser/Thr kinome. Hierarchichal clustering of 303 Ser/Thr kinase position specific scoring matrices (PSSMs). Kinase names are color-labeled according to their phylogenetic relationships (top right)^2^.

Overall inspection of sequence motifs associated with various branches of the motif-based dendrogram revealed that approximately ~60% of the Ser-Thr kinome could be represented by simple assignment to one of three previously observed motif classes: selectivity for basic residues N-terminal to the phosphorylation site (Cluster 1, Fig. 2), directed by a proline residue at the +1 position (Cluster 2), or a general preference for negatively charged (acidic and phosphorylated) residues at multiple positions (Cluster 3)^17,20,24,25^. Notably, more than half of all reported phosphorylation sites observed by MS could be assigned to one of these three signatures (Fig. S2). However, each of these motif classes could be further subcategorized based on selectivity both for and against distinct sets of residues at other positions, reflecting considerable diversity within these clusters (Figs. S3, S4, and S5).

The remaining ~40% of the Ser/Thr kinome comprised many smaller groups that displayed unique sequence determinants (Fig. 2, Clusters 4 – 17). For example, motifs for the DNA-damage response kinases (ATM, ATR, DNAPK and SMG1) clustered into a group that primarily selected a Gln residue at the +1 position (Cluster 5), consistent with previous studies^26,27^. Notably, several clusters displayed shared consensus motifs that have not been well recognized previously, such as the group including the IRAK, IRE, WNK, SNRK, and RIP kinases (Cluster 13), whose substrate motifs contained basic residues both N- and C-terminal to the phosphorylation site with dominant selection for aromatic residues in the +3 position. As another example, the kinases LKB1, CAMKK, PINK1, and PBK (Cluster 14) primarily recognized hydrophobic residues N-terminal to the phosphorylation site in combination with selection for turn-promoting residues (Gly or Asn) in the +1 position. Structural modeling of kinase-peptide complexes revealed complementary features within the kinase catalytic clefts likely responsible for recognition of these motifs (Fig. S6a,b).

An important and less generally recognized feature that dominated the clustering was strong negative selection against either positively or negatively charged residues at distinct positions within a motif, suggesting that electrostatic filtering strongly influences kinase substrate selection throughout the kinome^28,29^. We identified additional classes of amino acids, such as hydrophobic residues, that are selected against by a variety of kinases. These trends suggest that substrate avoidance plays a fundamental role in dictating correct kinase-substrate interactions^30,31^.

Unexpectedly, we observed that many kinases (129 out of 303) selected either a phospho-Thr or a phospho-Tyr as the preferred amino acid in at least one position within the motif (Fig. S1). In addition to kinases whose dependence on phospho-priming was previously known [GSK3, casein kinase 1, and casein kinase 2 families^32,33^, Cluster 3], this phenomenon was particularly evident for the GRK and YANK family kinases (Fig. S5), both of which have complementary basic residues within their catalytic domains (Fig. S6c,d). Intriguingly, individual GRK family members showed unique and specific selection for the location of the phospho-Thr or phospho-Tyr residue within their substrate peptides. GRKs are best known for their role in desensitization of G-protein coupled receptors (GPCRs), where multisite phosphorylation induces binding of arrestin proteins to inhibit signaling^34,35^. Our findings suggest that the capacity for only seven GRKs to differentially regulate 800 distinct GPCRs likely involves a complex interplay between initial sequence-specific phosphopriming of GPCRs by other serine/threonine and tyrosine kinases, followed by a second level of specificity resulting from GRK-dependent phosphorylation and subsequent recognition by a small number of β-arrestins.

Features of substrate recognition motifs across the entire kinome could be structurally rationalized based on the presence of specificity-determining residues at particular positions within the kinase catalytic domain^36,37^, leading to both expected and unexpected discoveries. For example, we found half of the kinases to display some degree of selectivity for either a Ser or a Thr as the phospho-acceptor residue (Fig. S7). Consistent with our previously published observations^38^, Ser or Thr phospho-acceptor site preference strongly correlated with the identity of the ‘DFG+1’ residue within the kinase activation loop, with bulky residues (Phe, Trp, Tyr) at this position in Ser-selective protein kinases and β-branched residues (Val, Ile, Thr) at this position in Thr-selective kinases. For some DFG+1 residues, however, Ser vs. Thr selectivity was unexpectedly context dependent. For instance, a Leu residue in the DFG+1 position was observed in both Ser-selective and dual specificity kinases, while a DFG+1 Ala residue resulted in a preference for Thr phosphorylation in the context of some kinases (e.g., the mitogen-activated protein kinase kinase kinases [MAP3Ks]), but a preference for Ser specificity in others (the IκB kinases [IKKs]). These observations, notable only within the context of the complete Ser/Thr kinome, indicate that additional residues beyond the previously established DFG+1 position can influence Ser/Thr specificity in a context-dependent manner.

### Phosphorylation motifs for the entire human serine/threonine kinome allow comprehensive annotation of the human phosphoproteome

Comprehensive knowledge of the human Ser/Thr kinase specificity has the potential to ‘de-orphanize’ the large number of reported phosphorylation sites with no associated kinase. To do so we generated a kinome-wide annotation of the human Ser/Thr phosphoproteome by computationally ranking each of ~50,000 high confidence phosphorylation sites against each Ser/Thr kinase motif (Fig. 3a, Table S2)^39^. Interestingly, more than 98% of these phosphorylation sites ranked favorably for at least one kinase we profiled (i.e., the site scored in the top 10% of all sites in the human phosphoproteome for that kinase). These annotations were strongly concordant with sites for which protein kinases involved have been previously identified. For phosphorylation sites whose upstream kinase has been previously verified by at least 3 independent reports, encompassing 969 sites and over 1/3^rd^ of the kinome, our motif-based approach yielded a median percentile of 93% (i.e., the reported site received a higher score than 93% of all putative phosphorylation sites in the phosphoproteome for its established kinase) (Fig. S8a). Furthermore, when we back-mapped the motifs of all 303 profiled kinases onto the literature-reported phosphorylation sites, our approach yielded a median reported kinase percentile of 92%, (i.e., the reported kinase scored more favorably than 92% of all profiled kinases in our atlas for its established substrate) (Fig. S8b). These rankings further improved when we considered kinase-substrate pairs with higher numbers of prior reports (Figs. S9, S10), suggesting that in a large majority of cases the linear sequence context of phosphorylation sites contributes substantially to kinase-substrate relationships.

**Fig. 3.**
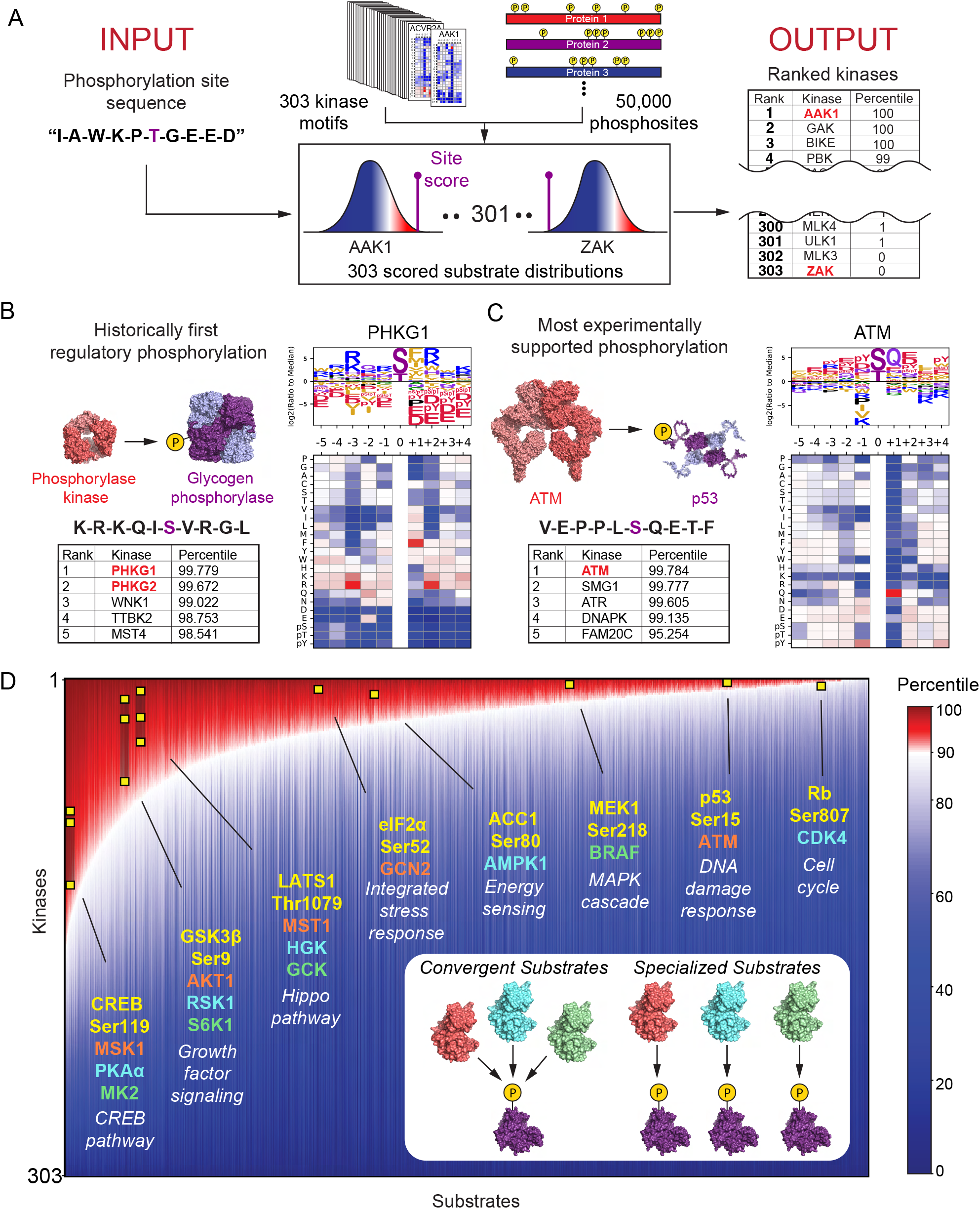
Phosphorylation motifs for the human serine/threonine kinome allow comprehensive scoring and annotation of the human phosphoproteome. (A) Schematic of the substrate scoring process. (B) Results for Ser15 on glycogen phosphorylase alongside PSSM and substrate motif logo of its established kinase glycogen phosphorylase kinase. (C) Results for Ser15 on p53 alongside its established kinase ATM. (D) Annotation of the human Ser and Thr phosphoproteome by percentile-scores from 303 Ser/Thr kinases as illustrated in (A). ~50,000 scored phosphorylation sites were sorted along the x-axis by their median kinase percentile-score. On the y-axis, kinase percentile scores were sorted by rank separately for each site and represented by heatmap. Examples of well-studied kinase-substrate relationships are highlighted (yellow squares). Inset: Phosphorylation sites on the left end of plot scored favorably for many kinases while sites on the right end scored favorably for fewer kinases.

Remarkably, motif predictions alone successfully identified numerous prominently studied kinase substrate relationships. For example, phosphorylase kinases PHKG1 and PHKG2 emerged as the top two hits (out of 303 kinases) for phosphorylating Ser15 of glycogen phosphorylase (Fig. 3b). This phosphoregulatory event, the very first to be discovered^40,41^, opened up the entire field of phosphorylation-dependent signal transduction. The most highly cited kinase-substrate interaction reported to date is phosphorylation of the tumor suppressor p53 at Ser15 by the DNA damage-activated kinase ATM, which scored as the top-ranking kinase associated with that site (Fig. 3c). Notably, other kinases reported to phosphorylate the same site, ATR, SMG1, and DNAPK, scored within the top 4 predicted kinases^42,43^.

Our approach could also correctly identify kinases for phosphorylation events driven by substrate co-localization or non-catalytic docking interactions, where we expected less dependence on the phosphorylation site motifs of their kinases. For example, we correctly identified both the mitochondrial-localized phosphorylation of pyruvate dehydrogenase by the pyruvate dehydrogenase kinases (Fig. S11a) and the docking-driven phosphorylation of the MAP kinase ERK by MEK (Fig. S11b)^44–48^. Interestingly, the phosphorylation site on ERK was selected *against* by nearly every human protein kinase we profiled except MEK, explaining how ERK can be exclusively regulated by MEK while avoiding phosphorylation by the kinome at large. Finally, our approach could tease apart kinase subfamilies with similar motifs and correctly assign them to their established substrates. For example, we could distinguish the CDK family kinases assuming classical roles in cell cycle progression (CDK1,2,3,4 and 6) from the subset of CDKs that govern gene transcription (CDK7,8,9,12,13 and 19) (Fig. S12)^49,50^.

Functional annotation of the human phosphoproteome allowed us to explore global trends in kinase-substrate interactions. We found that most phosphorylation sites could be assigned to a very small number of putative kinases (i.e., BRAF-MEK1, ATM-p53, and CDK4-Rb in Fig. 3d). However, a substantial minority of sites lacked unique negative sequence-discriminating features, and instead matched well to the optimal phosphorylation motifs for a greater number of kinases (i.e., Ser119 of CREB, Ser9 of GSK3B, and Thr1079 of LATS1; Fig. 3d) ^25,51–53^. This could suggest the importance of other kinase-determining factors (scaffolds, localization, etc.) for proper kinase-substrate recognition, or may indicate that these specific phosphorylation sites are points of convergence for multiple signaling pathways. For example, cAMP response element binding protein (CREB) is canonically phosphorylated at Ser119 by cAMP-dependent protein kinase (PKA), however, numerous prior reports demonstrate that a broad range of cellular stimuli and drug perturbations impinge on phosphorylation of this site by no less than ten distinct kinases^15,54^. Taken together, these findings suggest that the presence of negative selectivity elements flanking a putative phosphorylation site can be used to insulate a substrate from inappropriate phosphorylation by dozens of related kinases, while the absence of such negative selectivity can allow protein kinases in distinct pathways to converge on the same target.

### Global motif analysis reveals how kinase perturbations and pathway rewiring reshape the phosphoproteome

Cell signaling networks are complex and dynamic. Perturbation of kinase signaling pathways by genetic manipulations, treatment with growth factors and ligands, environmental stress, or small molecule inhibitors reshapes the phosphoproteome through both direct and indirect effects as a consequence of secondary signaling responses and/or off-target effects from the experimental treatment^13,55,56^. Due to the interconnected and dynamic nature of phosphorylation networks, distinguishing initial signaling events from those that result from the subsequent activation of additional signaling pathways is a common and challenging problem. We reasoned that kinases underlying both primary and secondary phosphorylation events could potentially be revealed by a global motif-based analysis of changes in the corresponding phosphoproteome. To test this idea, we used publicly available MS datasets from cells collected in the absence or presence of various perturbations and scored all phosphorylation sites with our atlas of serine/threonine kinase motifs. Kinase motifs significantly enriched or depleted following experimental treatment were then represented as volcano plots of motif frequencies and adjusted p-values (Fig. 4a).

**Fig. 4.**
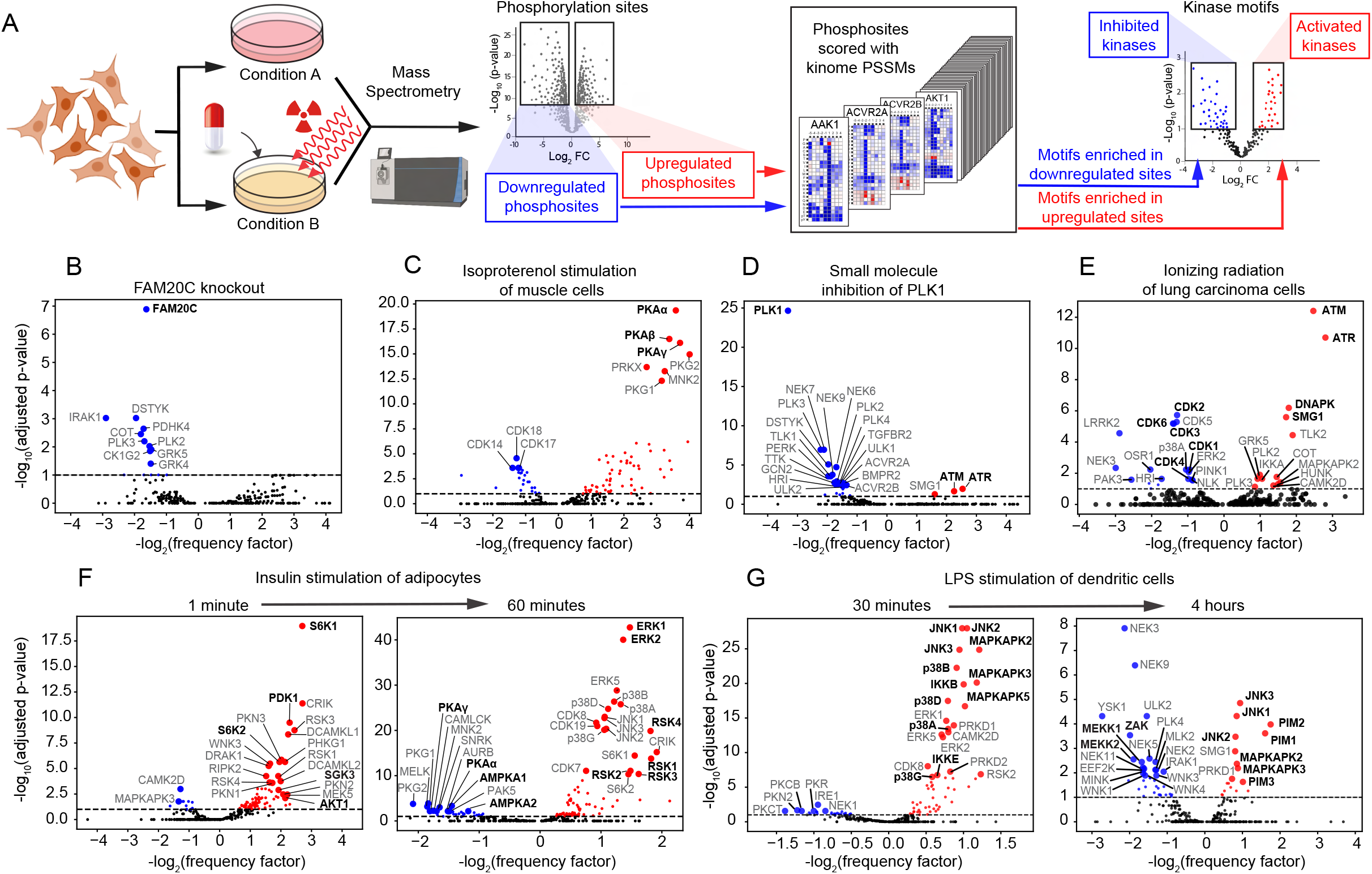
Global motif analysis reveals how kinase perturbations and pathway rewiring reshape the phosphoproteome. (A) Schematic depiction of motif enrichment analysis of phosphoproteomics data. (B-G) Results from published datasets: (B) conditioned medium of HepG2 cells following genetic deletion of FAM20C^58^, (C) cultured myotubes following 30-minute treatment with 2 μM isoproterenol^59^, (D) HeLa cells following mitotic arrest by 45-minute treatment with 0.1 μM PLK1 inhibitor BI-2536^62^, (E) A549 cells 2 hours following exposure to 6 Gy of ionizing radiation^65^, (F) 3T3-L1 adipocytes following serum starvation and then 1-minute and 60-minute treatment with 100 nM insulin^70^, (G) C57BL/6J mouse bone-marrow derived dendritic cells following 30-minute and 4-hour treatment with 100 ng/mL LPS^71^.

Using this approach, we found that sequence motifs corresponding to the most direct target of a genetic or chemical perturbation were among the most significantly regulated (seen, for example, with genetic deletion of the secreted primordial casein kinase FAM20C (Fig. 4b))^57^. When quantitative phosphoproteomics data from HepG2 cells lacking FAM20C^58^ were analyzed by our kinome-wide dataset, the most downregulated kinase recognition motif corresponded to that of FAM20C. Similarly, when skeletal muscle-like myotube cells were stimulated for 30 minutes with isoproterenol^59^, the most upregulated phosphorylation motifs corresponded to multiple isoforms of cAMP-dependent protein kinase (PKA), canonical effector kinases downstream of the β_1_ and β_2_ adrenergic receptors (Fig. 4c)^60,61^. Of note, PKA motifs are highly similar to those of several other basophilic kinases yet we could identify their enrichment in this scenario. In addition, our comprehensive serine/threonine kinome motif collection elucidated secondary signaling events in a dataset from HeLa cells arrested in mitosis using the PLK1 inhibitor BI-2536 (Fig. 4d)^62^ where, in addition to observing a striking downregulation of substrates containing the optimal PLK1 motif, we also noted significant upregulation of substrates phosphorylated by ATM and ATR. This finding is in good agreement with prior reports that PLK1 can suppress DNA damage signaling in mitotic cells^63,64^.

Our motif-based analysis could also be used to reveal key signaling events resulting from more complex interventions. For example, we interrogated phosphoproteomic data from A549 cells treated with 6 Gy of ionizing radiation (Fig. 4e)^65^. Our analysis revealed the up- and down-regulation of numerous signaling pathways, including upregulation of canonical kinases involved in the DNA damage response (ATM, ATR, DNAPK, SMG1) and downregulation of canonical kinases involved in cell cycle progression (CDK1, 2, 4, and 6) consistent with G1/S and G2/M arrest. Furthermore, we found up- and down-regulation of less appreciated DNA damage-responsive kinases [MAPKAPK2 ^66,67^, PLK3 ^68^, and LRKK2 ^69^].

The full collection of serine/threonine kinome motifs also allowed temporal dynamics of signaling to be resolved from time-resolved phosphoproteomic datasets. For example, motif-based analysis of phosphoproteomic data from insulin-treated 3T3-L1 adipocytes^70^ revealed rapid activation of the phosphoinositide 3-kinase signaling pathway within 1 minute after insulin stimulation followed by subsequent activation of the MAPK pathway, together with downregulation of AMP- and cAMP-dependent protein kinases within 60 minutes (Fig. 4f). Similarly, phosphoproteomic data analysis from LPS-stimulated dendritic cells^71^ suggested marked upregulation at 30 minutes of a set of stress-activated kinases including the IKKs, JNK and p38 MAPKs, along with the MAPKAPK family of p38 effector kinases, followed within 4 hours by subsequent upregulation of the PIM kinases and suppression of the MAPKs in parallel with the downregulation of their upstream MAPK3Ks (MEKK1, MEKK2, and ZAK)^72,73^, suggestive of a negative feedback loop (Fig. 4g). Thus, comprehensive motif-based approaches, when applied to time-resolved phosphoproteomics experiments, can decipher the distinct temporal dynamics of different groups of kinases.

## DISCUSSION

This work presents the full spectrum of substrate motifs of the human serine/threonine kinome and provides an unbiased global framework to further explore their cellular functions. Globally, these motifs are substantially more diverse than expected, suggesting a broader substrate repertoire of the kinome. Hierarchical clustering of this dataset reorganized the kinome into 17 motif-classes and introduced several novel shared motif features (Fig. 2). For kinases with similar motifs, we saw multiple cases where their minor differences translated into dramatically different substrate predictions and motif enrichments (Fig. S11, Fig. 4c-g) and rationalized how such biochemically similar kinases can have divergent biological roles.

The serine/threonine kinases we profiled were, almost without exception, strongly discriminatory against specific motif features. This negative selection rationalizes how kinases sharing similar positively selected residues in their motifs can regulate distinct signaling pathways with specialized cellular functions, and how substrates are insulated from inappropriate phosphorylation by irrelevant kinases. Intriguingly, these findings suggest that fidelity in kinase signaling pathways is largely achieved through selective pressure on substrates to avoid phosphorylation by the majority of irrelevant kinases, and that this may occur by tuning the amino acid sequences surrounding the phosphorylation sites to be disfavored by non-cognate kinases. Since this negative selection contributes substantially to proper substrate recognition, accurate identification of kinase-substrate relationships requires a comprehensive knowledge of kinase phosphorylation motifs – not only for an individual kinase of interest, but also for all other kinases in the human kinome that might compete for the same substrate pool.

When this kinome-wide dataset was used to predict specific kinases responsible for substrate phosphorylation solely based on the amino acid sequence surrounding the phosphorylation site, the results were remarkably accurate at identifying correct kinase-substrate relationships, even without knowledge of tissue specificity, scaffolding effects, or subcellular localization. Including such additional information is likely to further improve these predictive approaches^74,75^. Interrogation of MS phosphoproteomic datasets using this global collection of motifs yielded new potential biological insights and new putative kinase substrates (Fig. 4). For example, in cells undergoing exposure to ionizing radiation (Fig. 4e), ATM was predicted to target 35 of the phosphorylation sites that were upregulated, most of which have never been associated as substrates for ATM (Table S3). As the application of phosphoproteomics to human clinical samples and disease model systems continues to advance^76^, our comprehensive motif-based approach will be uniquely equipped to unravel complex signaling events that underlie human disease progressions, mechanisms of cancer drug resistance, dietary interventions, and other important physiological processes. In sum, we foresee this providing a valuable resource for a broad spectrum of researchers who study signaling pathways in human biology and disease.

## MATERIALS & METHODS

### Plasmids

For expression and purification from bacteria, DNA sequences for the human Ser/Thr kinases, kinase binding partners, and chaperones listed below were codon-optimized for E. coli using GeneSmart prediction software (Genscript). Optimized coding sequences were synthesized as gBlocks (Integrated DNA Technologies) carrying 16-base pair overhangs at the 5’ and 3’ ends to facilitate in-fusion cloning (Clontech) into pET expression vectors (EMD MIllipore).

#### pCDFDuet1 constructs

HSP90AA1-His_6_ (full length), hereafter referred to as “HSP90,” untagged HSP90 (full length), His_6_-MO25a (full length), His_6_-ALPHAK3/ALPK1 NTD (1-474), and His8-CCNC (full length) in tandem with MED12-His8 (1-100), and untagged CK2B (full length).

#### pET28a constructs

His_6_-PDPK1 (full length), His_6_-PRP4/PRPF4B (519-end), GST-CHAK1/TRPM6 (1699-end), His_6_-caMLCK/MYLK3 (490-end), untagged MEK5/MAP2K5-DD (full length), His_6_-ERK7/MAPK15 (full length), His_6_-SUMO-ALPHAK3/ALPK1 CTD (959-end), MYO3A-His_6_ (1-308), His_6_-NIK/MAP3K14 (327-673), and BMPR2-His_6_ (172-504).

#### pETDuet1 constructs

His_6_-CDK8 (full length), His_6_-CDK19 (full length), ERK5/MAPK7-His_6_ (1-405), His_6_-AAK1 (27-365), His_6_-BIKE (37-345), CK2A1-His_6_ (full length), CK2A2-His_6_ (full length), His_10_-MBP-MEKK1/MAP3K1 (1174-end), His_6_-CLK1 (128-end), His_8_-PLK2 (57-360), His_10_-MAP3K15 (631-922), His_6_-SUMO-ASK1/MAP3K5 (659-951), and His_6_-TAO2 (1-350).

#### pACYDuet1 construct

Untagged CDC37 (full length).

#### Mammalian expression constructs

For enhanced expression in mammalian lines cells, the DNA sequences of His_6_-GST-SBK (full length) and Flag-His_6_-WNK3 (1-434) were optimized for expression in H. sapiens using GeneSmart (Genscript) and synthesized as gBlocks (Integrated DNA Technologies) carrying 16-base pair overhangs to facilitate in-fusion cloning into digested pCDNA3.4 (Thermo).

To generate a mammalian expression construct for the TAK1/MAP3K7, the coding sequence for this kinase (GE Healthcare Dharmacon: MHS6278-202756930) and its binding partner TAB1 (GE Healthcare Dharmacon: MHS6278-202760135) were PCR amplified and ligated as a fusion construct (TAK1 (1-303)-TAB1(451-end)) into the mammalian expression vector pLenti-X by infusion.

#### Expression constructs purchased or obtained from other laboratories

Bacterial expression constructs for GST-VRK1 (full length) and GST-VRK2 (full length), in pGEX-4T, were received as gifts from Pedro Lazo at CSIC-Universidad de Salamanca ^1^. Bacterial expression construct for mouse CDKL5-His_6_ (1-352), in pET23a+, was received as a gift from Syouichi Katayama at Ritsumeikan University ^2^. Bacterial expression constructs for His_6_-SUMO-PDHK1 (full length), His_6_-SUMO-PDHK4 (full length), pGroESL (GroEL/GroES), and MBP-BCKDK (full length) were received as gifts from David Chuang, Shih-Chia Tso, and Richard Wynn at UT Southwestern Medical Center ^3,4^. pProEx HTa-BRAF_16mut V600E (444-721) was a gift from Marc Therrien at Université de Montréal ^5^. Mammalian expression constructs for Flag-ATR (S1333A) and HA-ATRIP were provided by David Cortez at Vanderbilt University School of Medicine ^6^. Bacterial expression constructs for DMPK1, CAMK1A, CAMK1G, CAMK2G, PHKG2, CDKL1, GAK, and lambda phosphatase were purchased from Addgene (Addgene Kit #1000000094)^7^.

### Expression and Purification from bacteria

Transformations were performed with BL21 Star cells (Thermo Fisher) unless specified otherwise. Antibiotic concentrations used: Carbenicillin (100 mg/L), Kanamycin (50 mg/L), Spectinomycin (25 mg/L), and Chloramphenicol (25 mg/L in EtOH, prepared fresh). Transformed cells were grown in 1L Terrific broth by shaking at 190 rpm at 37°C until optical density reached 0.7-0.8, at which point 1mM IPTG was added to induce expression. The cells were then transferred to a refrigerated shaker and shaken at 220 rpm at 18°C for 16-20 hours. Cells were centrifuged at 6,000 x g, and pellets were snap freezed in liquid nitrogen and stored at −80 °C.

All steps in the protein purification were performed at 4°C. Cell pellet was solubilized in lysis buffer (see contents below), using spatula to disperse, and lysed by probe sonication. The lysate was centrifuged at 20,000 x g for 1 h and the supernatant was combined with affinity purification resin, nickel NTA (Qiagen) or glutathione sepharose (GE Health), that had been rinsed in base buffer. The supernatant-bead slurry was agitated using a rotisserie for 30 minutes. Resin was washed with 1 L base buffer and eluted in 10 bed volumes of elution buffer. Eluted protein was concentrated using Ultra Centrifugal Filter Units (Amicon), supplemented with 1 mM DTT and 25% glycerol, and snap freezed in liquid nitrogen and stored at −80 °C.

Standard lysis buffer: 50 mM Tris pH 8.0, 100 mM NaCl, 2 mM MgCl2, 2% glycerol, HALT EDTA-free phosphatase and protease inhibitor cocktail (Life technologies), 5 mM betamercaptoethanol, 1-3 grams of lysozyme (Sigma)

Standard base buffer: 50 mM Tris pH 8.0, 100 mM NaCl, 2 mM MgCl2, 2% glycerol (include 50 mM imidazole for purifications involving polyhistidine-tag)

Standard wash buffer: 50 mM Tris pH 8.0, 500 mM NaCl, 2 mM MgCl2, 2% glycerol (include 50 mM imidazole for purifications involving polyhistidine-tag)

Polyhistidine-tag elution buffer: 50 mM Tris pH 8.0, 100 mM NaCl, 2 mM MgCl2, 2% glycerol, 350 mM imidazole

GST-tag elution buffer: 50 mM Tris pH 8.0, 100 mM NaCl, 2 mM MgCl2, 2% glycerol, 10 mM glutathione (pH adjusted after addition of glutathione)

The kinases BRAF and NIK were co-expressed with untagged HSP90/CDC37. CDK8 was copurified with CCNC/MED12. CDK19 was co-purified with CCNC/MED12 pCDFDuet1. CK2A1 and CK2A2 were co-purified with CK2B. ERK5 was co-expressed with MEK5DD. ALPHAK3 NTD (pCDFDuet1) and CTD (pETDuet1) were co-purified. DMPK1, CAMK1A, CAMK1G, CAMK2G, PHKG2, CDKL1, and GAK were co-expressed with lambda phosphatase in Rosetta 2 cells (Novagen).

PDHK1, PDHK4, BCKDK were co-expressed with GroeL/GroeS and purified with the following buffers: Lysis buffer (100 mM potassium phosphate pH 7.5, 10 mM L-arginine (stock pH-adjusted to 7.5), 500 mM KCl, 0.1 mM EDTA, 0.1 mM EGTA, 0.2% Triton X-100, lysozyme), wash buffer (50 mM potassium phosphate pH 7.5, 10 mM arginine, 500 mM NaCl, 0.1% Triton X-100, 2 mM MgCl2), and Elution buffer (25 mM Tris pH 7.5, 120 mM KCl, 0.02% Tween-20, 50 mM Arginine, 350 mM imidazole for PDHK1 and PDHK4 only, 20 mM maltose for BCKDK only). BCKDK was purified by its MBP tag on amylose resin (NEB).

CDKL5 was expressed in BL21-codonplus(DE3)-RIL cells.

KIS (full length) was purified as described previously^8^.

### Expression and purification from mammalian cells

#### Transfection

Expi293F cells (Thermo Fisher) were cultured in 500mL Expi293 Expression Medium (Thermo Fisher) in 2L spinner flasks on a magnetic stirring platform at 100 RPM at 36.8°C with 8% CO2. For transfection, 500 μg of expression constructs were diluted in Opti-MEM I Reduced Serum Medium (Thermo Fisher). ExpiFectamine 293 Reagent (Thermo Fisher) was diluted with Opti-MEM separately then combined with diluted plasmid DNA for 10 minutes at room temperature. The mixture was then transferred to the cells (3 X 10^6^ cells/mL) and stirred. 20 hours after transfection, ExpiFectamine 293 Transfection Enhancer 1 and Enhancer 2 (Thermo Fisher) were added to the cells. Two days later, the cells were centrifuged at 300 X g for 5 min, snap freezed in liquid nitrogen, and stored at −80°C (3 days post-transfection).

#### Purification

All steps of protein purification were performed at 4°C. Cell pellet was solubilized in lysis buffer, using spatula to disperse, and lysed by Dounce homogenization (20 strokes). The lysate was centrifuged at 100,000 x g for 1 h and the supernatant was combined with affinity purification resin, nickel NTA (Qiagen), glutathione sepharose (GE Health), or Anti-Flag M2 affinity gel (Sigma), and agitated on rotisserie for 30 min (nickel and glutathione beads) or 1 hour (Anti-flag beads). Resin was washed with 1 L base buffer and eluted in 10 bed volumes of elution buffer. For elution of flag tagged-proteins, beads were immersed in elution buffer (0.15 ug/mL 3X Flag peptide (Sigma)) and agitated on rotisserie for 1 hour prior to elution. Eluted protein was concentrated using Ultra Centrifugal Filter Units (Amicon), supplemented with 1 mM DTT and 25% glycerol, and snap freezed in liquid nitrogen and stored at −80°C.

Standard lysis buffer: 50 mM Tris pH 8.0, 150 mM NaCl, 2 mM MgCl2, 5% glycerol, 1% Triton X-100, 5 mM β-mercaptoethanol, HALT protease inhibitors.

Standard base buffer: 50 mM Tris pH 8.0, 100 mM NaCl, 2 mM MgCl2, 2% glycerol.

Standard wash buffer: 50 mM Tris pH 8.0, 500 mM NaCl, 2 mM MgCl2, 2% glycerol.

His_6_-GST-tagged SBK was purified sequentially on nickel and then glutathione resins. The first wash buffer included 25 mM imidazole. SBK1 elution buffer for polyhistidine tag: 50 mM Tris pH 8.0, 100 mM NaCl, 2 mM MgCl2, 2% glycerol, 250 mM imidazole. SBK1 elution buffer for GST tag: 50 mM Tris pH 8.0, 100 mM NaCl, 2 mM MgCl2, 2% glycerol, 10 mM glutathione.

Flag-TAK1-TAB1 elution buffer: 50 mM Tris pH 8.0, 100 mM NaCl, 2 mM MgCl2, 2% glycerol, 0.15 ug/mL 3X Flag peptide.

Flag-His_6_-WNK3 was purified sequentially on nickel and then anti-flag resins. The first wash buffer contained 25 mM imidazole. Flag-tag elution buffer (chloride-free): 50 mM Tris pH 7.5, 2 mM MgAc2, 2% glycerol, 0.15 ug/mL 3X Flag peptide.

350 uL Flag-ATR (S1333A) and 150 ug Ha-ATRIP were co-transfected into Expi293 cells and incubated for one additional day following addition of enhancers (4 days post-transfection).

ATR lysis buffer: 50 mM HEPES pH 7.4, 150 mM NaCl, 10% glycerol, 0.25% Tween 20, 2 mM MgCl2, DTT.

ATR wash buffer: 50 mM HEPES pH 7.4, 150 mM NaCl, 0.01% Brij-35, 2 mM MgCl2, 5 mM ATP, DTT.

ATR elution buffer: 20 mM HEPES pH 7.4, 150 mM NaCl, 0.01% Brij-35, DTT, 0.15 ug/mL 3X Flag peptide.

Eluates were concentrated to 1 mL in 100K MWCO Amicon tubes and resolved by MonoS column in in 0-1M NaCl gradient (buffer 25 mM Bis Tris pH 6.9, 0.01% Brij-35, and 5 mM TCEP). 1 mL fraction were collected. Fractions 1-4 were combined and concentrated to 1 mL in 100K MWCO and resolved by size exclusion (Superose 6) in 20 mM HEPES pH 7.4, 200 mM NaCl, 0.01% Brij-35, and 5 mM TCEP. 1 mL fraction were collected. Fractions 11-14 were verified to be pure ATR:ATRIP complex on SDS-PAGE and profiled in PSPA.

SMG1:SMG9 complexes were purified from HEK293T cells as described previously^9^.

RIPK1, RIPK2, and RIPK3 were purified from insect cells as described previously^10^.

#### Recombinant active kinases obtained from other laboratories

Recombinant active CDK12:CycK, CDK13:CycK, and CDK9:CycT complexes were provided as gifts from Matthias Geyer at University of Bonn^11,12^.

Recombinant active DCAMKL1/DCLK1 and MELK were provided as gifts from Nathanael Gray and Kenneth Westover^13,14^.

Recombinant active PRPK(full length):CGI121(full length) complex was provided as a gift from Leo Wan and Frank Sicheri at the Lunenfeld-Tanenbaum Research Institute^15^.

Recombinant active HASPIN (452-798) was provided as a gift from Andrea Musacchio at the Institute of Molecular Physiology in Dortmund^16^.

Recombinant active YSK1 was provided as a gift from Xuelian Luo at UT Southwestern Medical Center^17^.

Catalog and lot numbers of purchased recombinant kinases are listed in Table S1.

### Peptide library arrays

Recombinant kinase was distributed across 384-well plate, mixed with the peptide substrate library in solution phase (Anaspec #AS-62017-1 and #AS-62335), and incubated for 90 mins (Assay conditions for each kinase described in Table S1)^18–22^. Each well contains a mixture of peptides with a centralized phosphor-acceptor (serine and threonine at a 1:1 ratio) and one fixed amino acid in a randomized background. All 20 natural amino acids, plus two PTM residues (phospho-Thr and phospho-Tyr), were substituted into positions −5 to +4 to generate 198 unique peptide mixtures (22 amino acids X 9 fixed positions). After reaction, peptides were separated were spotted on streptavidin-conjugated membranes (Promega #V2861) where they tightly associated via their C-terminal biotinylations. The membranes were rinsed to remove free ATP and kinase and imaged with Typhoon FLA 7000 phosphorimager (GE). Raw data (GEL file) was quantified by ImageQuant (GE). For the kinase ALPHAK3, spots were normalized to surrounding background, due to spatial variation in background signal. PDHK1 and PDHK4 showed dual specificity for serine and tyrosine. For these kinases, we utilized a customized peptide substrate library devoid of tyrosines at randomized positions.

In total, 286 human kinase motifs, one motif from a mouse kinase ortholog (CDKL5), and one motif from an arthropod Pediculus humanus corporis kinase ortholog (PINK1), were combined with 15 human kinase motifs we previously published, that included AKT1^23^, SRPK1^24^, SRPK2^24^, SRPK3^24^, CK1D^24^, DYRK1A^25^, DYRK2^25^, GSK3A^25^, GSK3B^25^, CK1A^25^, CK1E^25^, CK1G1^25^, CDK10^26^, CDK2^27^, CDK3^27^, CDK18^27^, and CDK7^28^.

### Matrix processing

The matrices were column-normalized (at all positions) by the sum of the 17 randomized amino acids (excluding serine, threonine, and cysteine), to yield positional specific scoring matrices (PSSMs). PDHK1 and PDHK4 were normalized by the 16 randomized amino acids (excluding serine, threonine, cysteine, and additionally tyrosine), corresponding to the uniquely customized peptide library that profiled these kinases. The cysteine row was scaled by its median to be 1/17 (1/16 for PDHK1 and PDHK4). The serine and threonine values in each position were set to be the median of that position. The ratio of serine vs threonine phospho-acceptor favorability (S_0_ and T_0_, respectively) was determined by summing up the values of the serine and threonine rows in the densitometry matrix (S_S_ and S_T_, respectively), and then normalized by the higher value among the two:

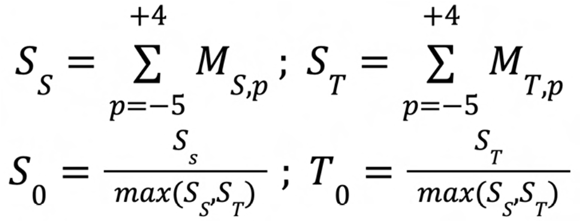

### Matrix clustering

The dendrogram in Fig. 2 was generated using the normalized matrices with the 20 unmodified amino acids, as well as phosphothreonine and phosphotyrosine. Linkage matrix was computed through the SciPy package in Python (v1.7.1), using ‘ward’ method. Results were converted to Newick tree format and plotted using FigTree (v1.4.4).

### Substrate scoring

For scoring substrates, the values of the corresponding amino acids in the corresponding positions were multiplied and scaled by the probability of a random peptide:

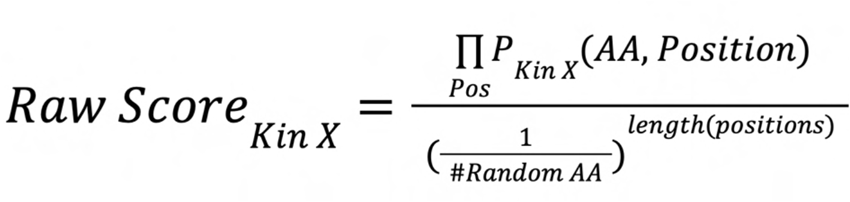

For the percentile-score of a substrate by a given kinase, we first computed the *a priori* score distribution of that kinase by scoring all the reported S/T phosphorylation sites on PhosphoSitePlus^29^ (downloaded on July 2021) with at least five high-throughput detections or one low-throughout detection, by the method discussed earlier. The percentile-score of a kinase-substrate pair is defined as the percentile ranking of the substrate within the score distribution of each kinase. This value was used for kinase enrichment as described before.

### Kinase enrichment analysis

The single phosphorylation sites (not including multi-phosphorylated peptides) in the analyzed phosphoproteomics studies were scored by all the characterized kinases (303 S/T kinases), and their ranks in the known phosphoproteome score distribution were determined as described above. For every non-duplicate, singly phosphorylated site, kinases that ranked within the top-15 kinases for the S/T kinases were considered as biochemically favored kinases for that phosphorylation site. For assessing kinase motif enrichment in phosphoproteomics datasets, we compared the percentage of phosphorylation sites for which each kinase was predicted among the upregulated/downregulated (increased/decreased, respectively) phosphorylation sites (sites with |log2Fold-Change| equal or greater than the logFC threshold), versus the percentage of biochemically favored phosphorylation sites for that kinase within the set of unregulated (unchanged) sites in this study (sites with |log2Fold-Change| less than the logFC threshold). LogFC threshold was determined to be 1.5 for all panels in Fig. 4, except for Fig. 4e where the threshold was set to 0.5 due to low range of the logFC in the data. Contingency tables were corrected using Haldane correction (adding 0.5 to the cases with zero in one of the counts). Statistical significance was determined using one-sided Fisher’s exact test, and the corresponding p-values were adjusted using the Benjamini-Hochberg procedure. Kinases that were significant (adjusted p-value <= 0.1) for both upregulated and downregulated analysis were excluded from downstream analysis. Then, for every kinase, the most significant enrichment side (upregulated or downregulated) was selected based on the adjusted p-value and presented in the volcano plots.

### Sequence logos

Sequence logos were made using logomaker package in Python^30^. For individual kinases, the normalized matrix was used, where the height of every letter is the ratio of its value to the median value of that position. The serine and threonine heights in the central position (position zero) were set to the ratio between their favorability, and to sum up to the maximal height in the peripheral positions. For clustered groups of kinases, the average matrix was calculated and presented as sequence logo as described above.

### Comparative analyses between amino acids in the kinase domains and their substrate specificities

For Fig. S7, kinases were sorted by their log2(S0/T0) values. For the sequence logo, kinase domains of 290 available kinases were obtained from previously aligned kinase sequences [PMID: 31875044]. The alignments to residues Met1-Leu296 in CDK2 (PDB: 1QMZ) were obtained for each kinase, and the frequencies of amino acids for each 15 kinases were calculated and plotted as a sequence logo.

### Known kinase-substrate pairs

Experimentally validated kinase-substrate relationships were obtained from PhosphoSitePlus (July 2021, Table S2). Number of reports for each pair was determined by the sum of the *in vivo* and *in vitro* reports.

### Illustrations

Experimental schema and illustrative models were generated by BioRender (https://biorender.com/) and Chemdraw. Kinome images generated and modified using Coral: http://phanstiel-lab.med.unc.edu/CORAL/. Structural illustrations were generated with PYMOL. Generic kinase domains in Figs 1 and 3: PKAα (pdb 1ATP). Kinase and substrate structures in Fig. 3: ATM (pdb 7SIC), p53 (chimera of alphaFold AF-P04637-F1-model_v2_1 (1-95) and 2ATA (96-292)), PHKG2 (pdb 2Y7J), and PYGM (pdb 1ABB)

## REFERENCES

1 Cohen, P. The origins of protein phosphorylation. Nature cell biology 4, E127–E130 (2002).

2 Manning, G., Whyte, D. B., Martinez, R., Hunter, T. & Sudarsanam, S. The protein kinase complement of the human genome. Science 298, 1912–1934 (2002).

3 Fuhs, S. R. & Hunter, T. pHisphorylation: the emergence of histidine phosphorylation as a reversible regulatory modification. Current opinion in cell biology 45, 8–16 (2017).

4 Hunter, T. Why nature chose phosphate to modify proteins. Philosophical Transactions of the Royal Society B: Biological Sciences 367, 2513–2516 (2012).

5 Blume-Jensen, P. & Hunter, T. Oncogenic kinase signalling. Nature 411, 355–365 (2001).

6 Lahiry, P., Torkamani, A., Schork, N. J. & Hegele, R. A. Kinase mutations in human disease: interpreting genotype–phenotype relationships. Nature Reviews Genetics 11, 60–74 (2010).

7 Ferguson, F. M. & Gray, N. S. Kinase inhibitors: the road ahead. Nature reviews Drug discovery 17, 353–377 (2018).

8 Wu, P., Nielsen, T. E. & Clausen, M. H. Small-molecule kinase inhibitors: an analysis of FDA-approved drugs. Drug discovery today 21, 5–10 (2016).

9 Berginski, M. E. et al. The Dark Kinase Knowledgebase: an online compendium of knowledge and experimental results of understudied kinases. Nucleic acids research 49, D529–D535 (2021).

10 Moret, N. et al. A resource for exploring the understudied human kinome for research and therapeutic opportunities. BioRxiv, 2020.2004. 2002.022277 (2021).

11 Needham, E. J., Parker, B. L., Burykin, T., James, D. E. & Humphrey, S. J. Illuminating the dark phosphoproteome. Science signaling 12, eaau8645 (2019).

12 Lemeer, S. & Heck, A. J. The phosphoproteomics data explosion. Current opinion in chemical biology 13, 414–420 (2009).

13 Aebersold, R. & Mann, M. Mass-spectrometric exploration of proteome structure and function. Nature 537, 347–355 (2016).

14 Riley, N. M. & Coon, J. J. Phosphoproteomics in the age of rapid and deep proteome profiling. Analytical chemistry 88, 74–94 (2016).

15 Hornbeck, P. V. et al. 15 years of PhosphoSitePlus®: integrating post-translationally modified sites, disease variants and isoforms. Nucleic Acids Research 47, D433–D441 (2019).

16 Kemp, B. E., Graves, D. J., Benjamini, E. & Krebs, E. G. Role of multiple basic residues in determining the substrate specificity of cyclic AMP-dependent protein kinase. Journal of Biological Chemistry 252, 4888–4894 (1977).

17 Kemp, B. E. & Pearson, R. B. Protein kinase recognition sequence motifs. Trends in biochemical sciences 15, 342–346 (1990).

18 Marin, O., Meggio, F., Marchiori, F., Borin, G. & Pinna, L. A. Site specificity of casein kinase-2 (TS) from rat liver cytosol: A study with model peptide substrates. European journal of biochemistry 160, 239–244 (1986).

19 Clark-Lewis, I., Sanghera, J. S. & Pelech, S. Definition of a consensus sequence for peptide substrate recognition by p44mpk, the meiosis-activated myelin basic protein kinase. Journal of Biological Chemistry 266, 15180–15184 (1991).

20 Pinna, L. A. & Ruzzene, M. How do protein kinases recognize their substrates? Biochimica et Biophysica Acta (BBA)-Molecular Cell Research 1314, 191–225 (1996).

21 Meggio, F. & Pinna, L. A. One-thousand-and-one substrates of protein kinase CK2? The FASEB Journal 17, 349–368 (2003).

22 Songyang, Z. et al. Use of an oriented peptide library to determine the optimal substrates of protein kinases. Current biology 4, 973–982 (1994).

23 Hutti, J. E. et al. A rapid method for determining protein kinase phosphorylation specificity. Nature methods 1, 27–29 (2004).

24 Mok, J. et al. Deciphering protein kinase specificity through large-scale analysis of yeast phosphorylation site motifs. Science signaling 3, ra12–ra12 (2010).

25 Pearce, L. R., Komander, D. & Alessi, D. R. The nuts and bolts of AGC protein kinases. Nature reviews Molecular cell biology 11, 9–22 (2010).

26 Kim, S.-T., Lim, D.-S., Canman, C. E. & Kastan, M. B. Substrate specificities and identification of putative substrates of ATM kinase family members. Journal of Biological Chemistry 274, 37538–37543 (1999).

27 O’Neill, T. et al. Utilization of oriented peptide libraries to identify substrate motifs selected by ATM. Journal of Biological Chemistry 275, 22719–22727 (2000).

28 Shah, N. H. et al. An electrostatic selection mechanism controls sequential kinase signaling downstream of the T cell receptor. Elife 5, e20105 (2016).

29 Shah, N. H., Löbel, M., Weiss, A. & Kuriyan, J. Fine-tuning of substrate preferences of the Src-family kinase Lck revealed through a high-throughput specificity screen. Elife 7, e35190 (2018).

30 Zhu, G. et al. Exceptional disfavor for proline at the P+ 1 position among AGC and CAMK kinases establishes reciprocal specificity between them and the proline-directed kinases. Journal of Biological Chemistry 280, 10743–10748 (2005).

31 Alexander, J. et al. Spatial exclusivity combined with positive and negative selection of phosphorylation motifs is the basis for context-dependent mitotic signaling. Science signaling 4, ra42–ra42 (2011).

32 Fiol, C. J., Wang, A., Roeske, R. W. & Roach, P. J. Ordered multisite protein phosphorylation. Analysis of glycogen synthase kinase 3 action using model peptide substrates. Journal of Biological Chemistry 265, 6061–6065 (1990).

33 Flotow, H. et al. Phosphate groups as substrate determinants for casein kinase I action. Journal of Biological Chemistry 265, 14264–14269 (1990).

34 Reiter, E. & Lefkowitz, R. J. GRKs and β-arrestins: roles in receptor silencing, trafficking and signaling. Trends in endocrinology & metabolism 17, 159–165 (2006).

35 Moore, C. A., Milano, S. K. & Benovic, J. L. Regulation of receptor trafficking by GRKs and arrestins. Annu. Rev. Physiol. 69, 451–482 (2007).

36 Bradley, D. et al. Sequence and Structure-Based Analysis of Specificity Determinants in Eukaryotic Protein Kinases. Cell reports 34, 108602 (2021).

37 Taylor, S. S. & Kornev, A. P. Protein kinases: evolution of dynamic regulatory proteins. Trends in biochemical sciences 36, 65–77 (2011).

38 Chen, C. et al. Identification of a major determinant for serine-threonine kinase phosphoacceptor specificity. Molecular cell 53, 140–147 (2014).

39 Obenauer, J. C., Cantley, L. C. & Yaffe, M. B. Scansite 2.0: Proteome-wide prediction of cell signaling interactions using short sequence motifs. Nucleic acids research 31, 3635–3641 (2003).

40 Fischer, E. H., Graves, D. J., Crittenden, E. R. S. & Krebs, E. G. Structure of the site phosphorylated in the phosphorylase b to a reaction. Journal of Biological Chemistry 234, 1698–1704 (1959).

41 Wolf, D. P., Fischer, E. H. & Krebs, E. G. Amino acid sequence of the phosphorylated site in rabbit liver glycogen phosphorylase. Biochemistry 9, 1923–1929 (1970).

42 Lakin, N. D. & Jackson, S. P. Regulation of p53 in response to DNA damage. Oncogene 18, 7644–7655 (1999).

43 Gehen, S. C., Staversky, R. J., Bambara, R. A., Keng, P. C. & O’Reilly, M. A. hSMG-1 and ATM sequentially and independently regulate the G1 checkpoint during oxidative stress. Oncogene 27, 4065–4074 (2008).

44 Linn, T. C., Pettit, F. H. & Reed, L. J. α-Keto acid dehydrogenase complexes, X. Regulation of the activity of the pyruvate dehydrogenase complex from beef kidney mitochondria by phosphorylation and dephosphorylation. Proceedings of the National Academy of Sciences 62, 234–241 (1969).

45 Kholodenko, B. N., Hancock, J. F. & Kolch, W. Signalling ballet in space and time. Nature reviews Molecular cell biology 11, 414–426 (2010).

46 Ubersax, J. A. & Ferrell Jr, J. E. Mechanisms of specificity in protein phosphorylation. Nature reviews Molecular cell biology 8, 530–541 (2007).

47 Endicott, J. A., Noble, M. E. & Johnson, L. N. The structural basis for control of eukaryotic protein kinases. Annual review of biochemistry 81, 587–613 (2012).

48 Xu, B.-e., Wilsbacher, J. L., Collisson, T. & Cobb, M. H. The N-terminal ERK-binding site of MEK1 is required for efficient feedback phosphorylation by ERK2 in vitro and ERK activation in vivo. Journal of Biological Chemistry 274, 34029–34035 (1999).

49 Malumbres, M. et al. Cyclin-dependent kinases: a family portrait. Nature cell biology 11, 1275–1276 (2009).

50 Eick, D. & Geyer, M. The RNA polymerase II carboxy-terminal domain (CTD) code. Chemical reviews 113, 8456–8490 (2013).

51 Cohen, P. & Frame, S. The renaissance of GSK3. Nature reviews Molecular cell biology 2, 769–776 (2001).

52 Meng, Z. et al. MAP4K family kinases act in parallel to MST1/2 to activate LATS1/2 in the Hippo pathway. Nature communications 6, 1–13 (2015).

53 Doble, B. W. & Woodgett, J. R. GSK-3: tricks of the trade for a multi-tasking kinase. Journal of cell science 116, 1175–1186 (2003).

54 Johannessen, M., Delghandi, M. P. & Moens, U. What turns CREB on? Cellular signalling 16, 1211–1227 (2004).

55 Cravatt, B. F., Simon, G. M. & Yates Iii, J. R. The biological impact of mass-spectrometry-based proteomics. Nature 450, 991–1000 (2007).

56 Rigbolt, K. T. & Blagoev, B. in Seminars in cell & developmental biology. 863–871 (Elsevier).

57 Tagliabracci, V. S., Pinna, L. A. & Dixon, J. E. Secreted protein kinases. Trends in biochemical sciences 38, 121–130 (2013).

58 Tagliabracci, V. S. et al. A single kinase generates the majority of the secreted phosphoproteome. Cell 161, 1619–1632 (2015).

59 Needham, E. J. et al. Phosphoproteomics of acute cell stressors targeting exercise signaling networks reveal drug interactions regulating protein secretion. Cell reports 29, 1524–1538. e1526 (2019).

60 Walsh, D., Perkins, J. P. & Krebs, E. G. An adenosine 3’, 5’-monophosphate-dependant protein kinase from rabbit skeletal muscle. Journal of Biological Chemistry 243, 3763–3765 (1968).

61 Sutherland, E. W. & Rall, T. The relation of adenosine-3’, 5’-phosphate and phosphorylase to the actions of catecholamines and other hormones. Pharmacological Reviews 12, 265–299 (1960).

62 Kettenbach, A. N. et al. Quantitative phosphoproteomics identifies substrates and functional modules of Aurora and Polo-like kinase activities in mitotic cells. Science signaling 4, rs5–rs5 (2011).

63 van Vugt, M. A. et al. A mitotic phosphorylation feedback network connects Cdk1, Plk1, 53BP1, and Chk2 to inactivate the G2/M DNA damage checkpoint. PLoS biology 8, e1000287 (2010).

64 Macůrek, L. et al. Polo-like kinase-1 is activated by aurora A to promote checkpoint recovery. Nature 455, 119–123 (2008).

65 Winter, M. et al. Deciphering the acute cellular phosphoproteome response to irradiation with X-rays, protons and carbon ions. Molecular & Cellular Proteomics 16, 855–872 (2017).

66 Reinhardt, H. C., Aslanian, A. S., Lees, J. A. & Yaffe, M. B. p53-deficient cells rely on ATM-and ATR-mediated checkpoint signaling through the p38MAPK/MK2 pathway for survival after DNA damage. Cancer cell 11, 175–189 (2007).

67 Reinhardt, H. C. & Yaffe, M. B. Kinases that control the cell cycle in response to DNA damage: Chk1, Chk2, and MK2. Current opinion in cell biology 21, 245–255 (2009).

68 Xie, S. et al. Plk3 functionally links DNA damage to cell cycle arrest and apoptosis at least in part via the p53 pathway. Journal of Biological Chemistry 276, 43305–43312 (2001).

69 Gonzalez-Hunt, C. et al. Mitochondrial DNA damage as a potential biomarker of LRRK2 kinase activity in LRRK2 Parkinson’s disease. Scientific reports 10, 1–12 (2020).

70 Humphrey, S. J. et al. Dynamic adipocyte phosphoproteome reveals that Akt directly regulates mTORC2. Cell metabolism 17, 1009–1020 (2013).

71 Mertins, P. et al. An integrative framework reveals signaling-to-transcription events in toll-like receptor signaling. Cell reports 19, 2853–2866 (2017).

72 Johnson, G. L. & Lapadat, R. Mitogen-activated protein kinase pathways mediated by ERK, JNK, and p38 protein kinases. Science 298, 1911–1912 (2002).

73 Gallo, K. A. & Johnson, G. L. Mixed-lineage kinase control of JNK and p38 MAPK pathways. Nature reviews Molecular cell biology 3, 663–672 (2002).

74 Miller, C. J. & Turk, B. E. Homing in: mechanisms of substrate targeting by protein kinases. Trends in biochemical sciences 43, 380–394 (2018).

75 Linding, R. et al. Systematic discovery of in vivo phosphorylation networks. Cell 129, 1415–1426 (2007).

76 Zhu, Y., Aebersold, R., Mann, M. & Guo, T. SnapShot: Clinical proteomics. Cell 184, 4840–4840. e4841 (2021).

## References

1 Sanz-García, M. et al. Substrate profiling of human vaccinia-related kinases identifies coilin, a Cajal body nuclear protein, as a phosphorylation target with neurological implications. Journal of proteomics 75, 548–560 (2011).

2 Sekiguchi, M. et al. Identification of amphiphysin 1 as an endogenous substrate for CDKL5, a protein kinase associated with X-linked neurodevelopmental disorder. Archives of biochemistry and biophysics 535, 257–267 (2013).

3 Wynn, R. M., Davie, J. R., Cox, R. P. & Chuang, D. T. Chaperonins groEL and groES promote assembly of heterotetramers (alpha 2 beta 2) of mammalian mitochondrial branched-chain alpha-keto acid decarboxylase in Escherichia coli. Journal of Biological Chemistry 267, 12400–12403 (1992).

4 Song, J.-L., Li, J., Huang, Y.-S. & Chuang, D. T. Encapsulation of an 86-kDa assembly intermediate inside the cavities of GroEL and its single-ring variant SR1 by GroES. Journal of Biological Chemistry 278, 2515–2521 (2003).

5 Thevakumaran, N. et al. Crystal structure of a BRAF kinase domain monomer explains basis for allosteric regulation. Nature structural & molecular biology 22, 37–43 (2015).

6 Luzwick, J. W., Nam, E. A., Zhao, R. & Cortez, D. Mutation of serine 1333 in the ATR HEAT repeats creates a hyperactive kinase. PLoS One 9, e99397 (2014).

7 Albanese, S. K. et al. An open library of human kinase domain constructs for automated bacterial expression. Biochemistry 57, 4675–4689 (2018).

8 Manceau, V. et al. Major phosphorylation of SF1 on adjacent Ser-Pro motifs enhances interaction with U2AF65. The FEBS journal 273, 577–587 (2006).

9 Melero, R. et al. Structures of SMG1-UPFs complexes: SMG1 contributes to regulate UPF2-dependent activation of UPF1 in NMD. Structure 22, 1105–1119 (2014).

10 Najjar, M. et al. Structure guided design of potent and selective ponatinib-based hybrid inhibitors for RIPK1. Cell reports 10, 1850–1860 (2015).

11 Czudnochowski, N., Bösken, C. A. & Geyer, M. Serine-7 but not serine-5 phosphorylation primes RNA polymerase II CTD for P-TEFb recognition. Nature communications 3, 1–12 (2012).

12 Greifenberg, A. K. et al. Structural and functional analysis of the Cdk13/Cyclin K complex. Cell reports 14, 320–331 (2016).

13 Liu, Y. et al. Chemical biology toolkit for DCLK1 reveals connection to RNA processing. Cell chemical biology 27, 1229–1240. e1224 (2020).

14 Ferguson, F. M. et al. Discovery of a selective inhibitor of doublecortin like kinase 1. Nature chemical biology 16, 635–643 (2020).

15 Beenstock, J. et al. A substrate binding model for the KEOPS tRNA modifying complex. Nature communications 11, 1–17 (2020).

16 Villa, F. et al. Crystal structure of the catalytic domain of Haspin, an atypical kinase implicated in chromatin organization. Proceedings of the National Academy of Sciences 106, 20204–20209 (2009).

17 Bae, S. J., Ni, L. & Luo, X. STK25 suppresses Hippo signaling by regulating SAV1-STRIPAK antagonism. Elife 9, e54863 (2020).

18 Murillo-de-Ozores, A. R., Chávez-Canales, M., de Los Heros, P., Gamba, G. & Castañeda-Bueno, M. Physiological processes modulated by the chloride-sensitive WNK-SPAK/OSR1 kinase signaling pathway and the cation-coupled chloride cotransporters. Frontiers in Physiology, 1353 (2020).

19 Filippi, B. M. et al. MO25 is a master regulator of SPAK/OSR1 and MST3/MST4/YSK1 protein kinases. The EMBO journal 30, 1730–1741 (2011).

20 Zhou, P. et al. Alpha-kinase 1 is a cytosolic innate immune receptor for bacterial ADP-heptose. Nature 561, 122–126 (2018).

21 Taipale, M. et al. Quantitative analysis of HSP90-client interactions reveals principles of substrate recognition. Cell 150, 987–1001 (2012).

22 Klatt, F. et al. A precisely positioned MED12 activation helix stimulates CDK8 kinase activity. Proceedings of the National Academy of Sciences 117, 2894–2905 (2020).

23 Balasuriya, N. et al. Phosphorylation-dependent substrate selectivity of protein kinase B (AKT1). Journal of Biological Chemistry 295, 8120–8134 (2020).

24 Yaron, T. M. et al. The FDA-approved drug Alectinib compromises SARS-CoV-2 nucleocapsid phosphorylation and inhibits viral infection in vitro. BioRxiv (2020).

25 Zheng, Y. et al. Regulation of folate and methionine metabolism by multisite phosphorylation of human methylenetetrahydrofolate reductase. Scientific reports 9, 111 (2019).

26 Robert, T. et al. Development of a CDK10/CycM in vitro kinase screening assay and identification of first small-molecule inhibitors. Frontiers in chemistry 8, 147 (2020).

27 Ferguson, F. M. et al. Discovery of covalent CDK14 inhibitors with pan-TAIRE family specificity. Cell chemical biology 26, 804–817. e812 (2019).

28 Rimel, J. K. et al. Selective inhibition of CDK7 reveals high-confidence targets and new models for TFIIH function in transcription. Genes & development 34, 1452–1473 (2020).

29 Hornbeck, P. V. et al. 15 years of PhosphoSitePlus®: integrating post-translationally modified sites, disease variants and isoforms. Nucleic Acids Research 47, D433–D441 (2019).

30 Wagih, O. ggseqlogo: a versatile R package for drawing sequence logos. Bioinformatics 33, 3645–3647 (2017).

