## Supplemental Figures for "A global atlas of substrate specificities for the human serine/threonine kinome"

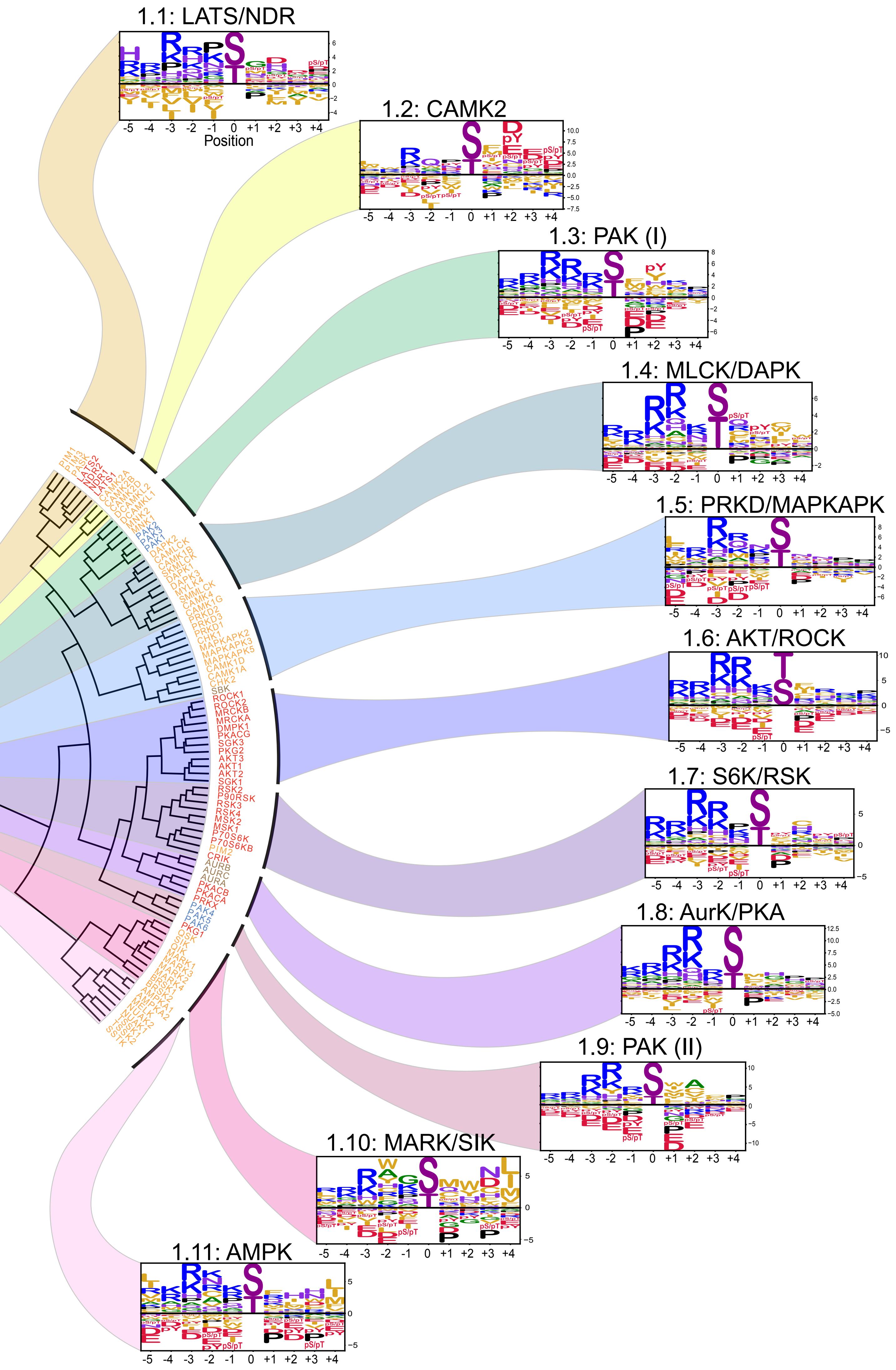


**Fig. S2 Subcategorization of the basophilic kinases of Cluster 1.**
Subcategorization of Cluster 1 from Figure 2 into 11 motif classes.

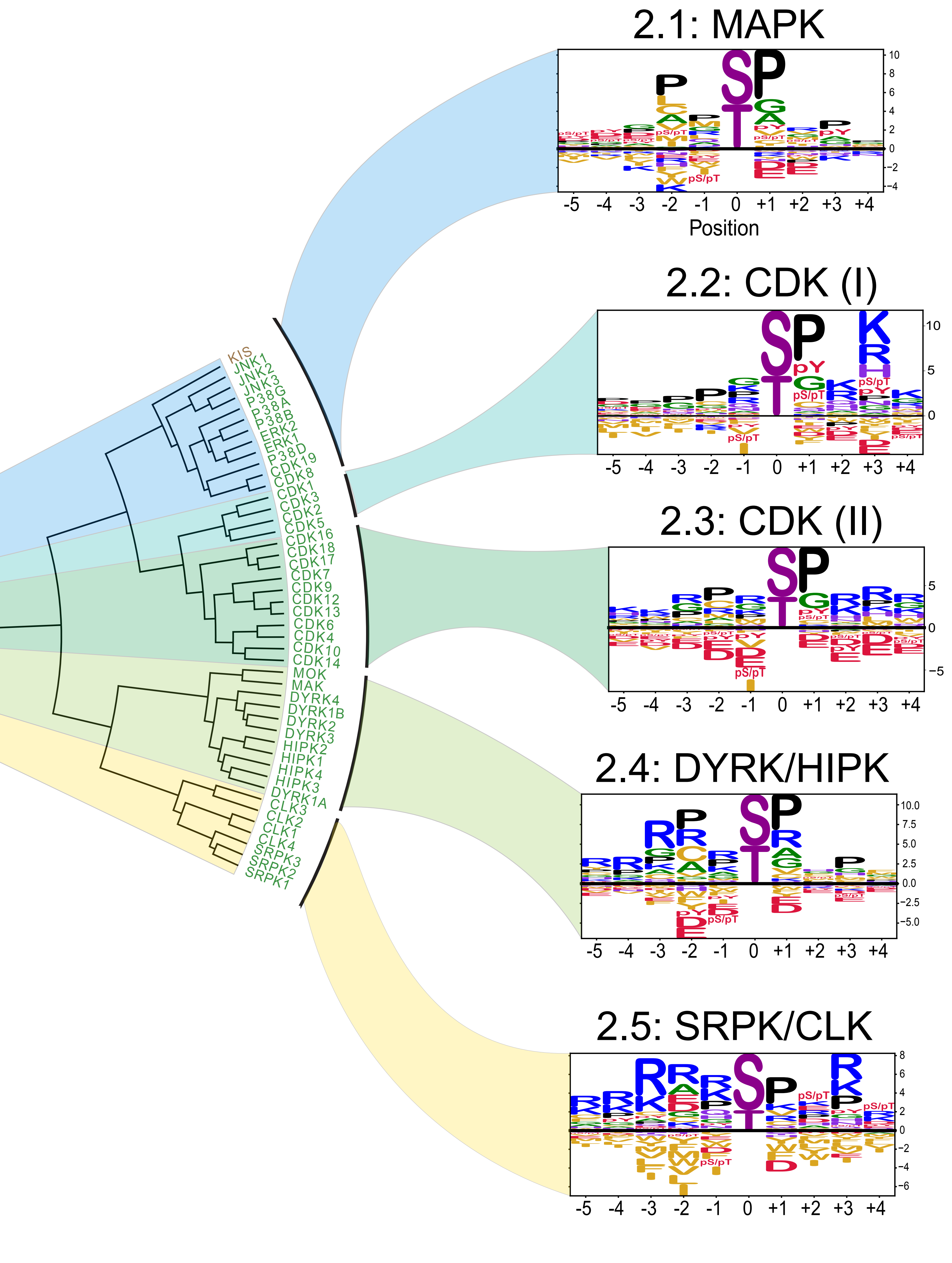


**Fig. S3 Subcategorization of the proline-directed kinases of Cluster 2.**
Subcategorization of Cluster 2 from Figure 2 into 5 motif classes.

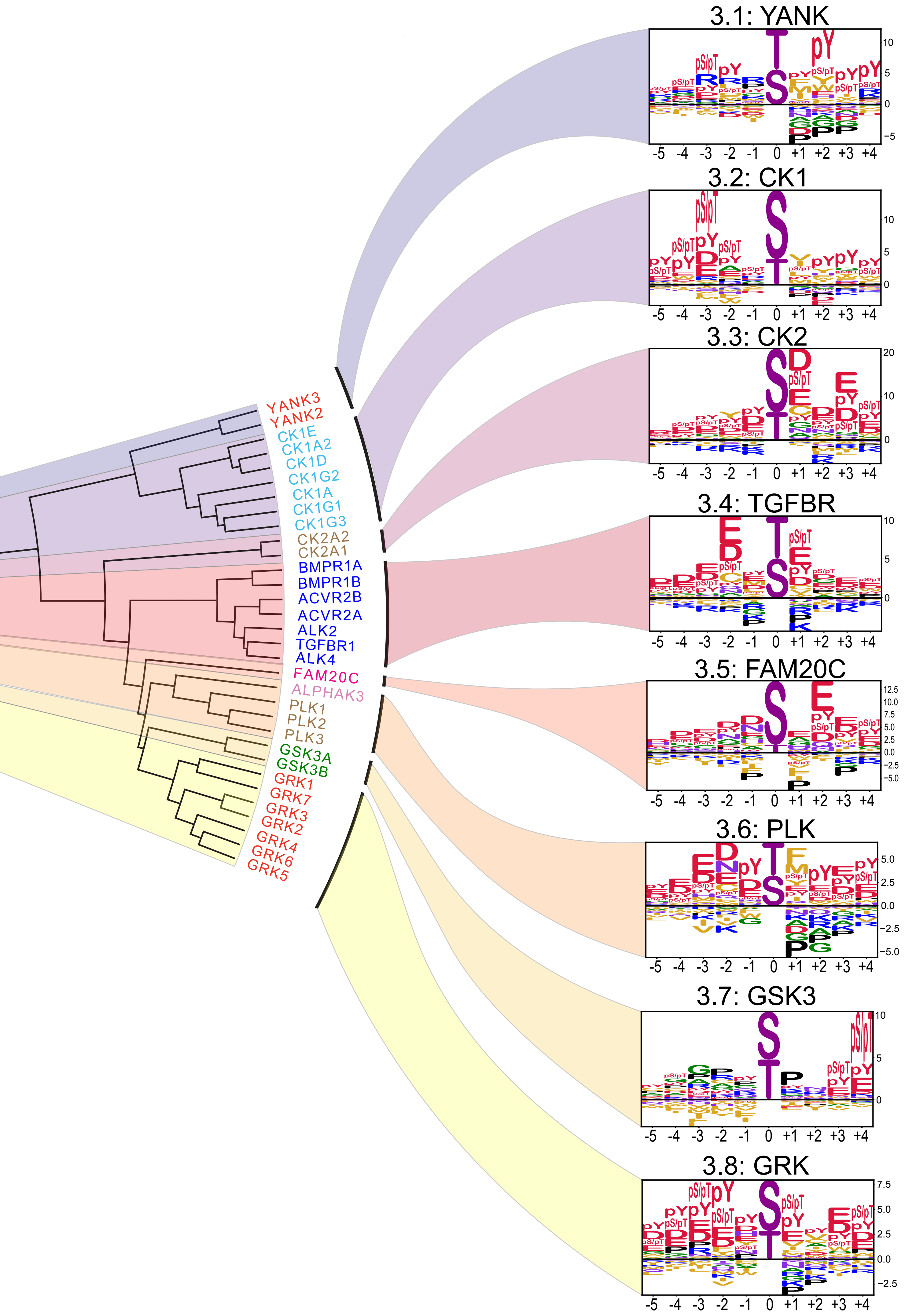


**Fig. S4 Subcategorization of the acidophilic kinases of Cluster 3.**
Subcategorization of Cluster 3 from Figure 2 into 8 motif classes.

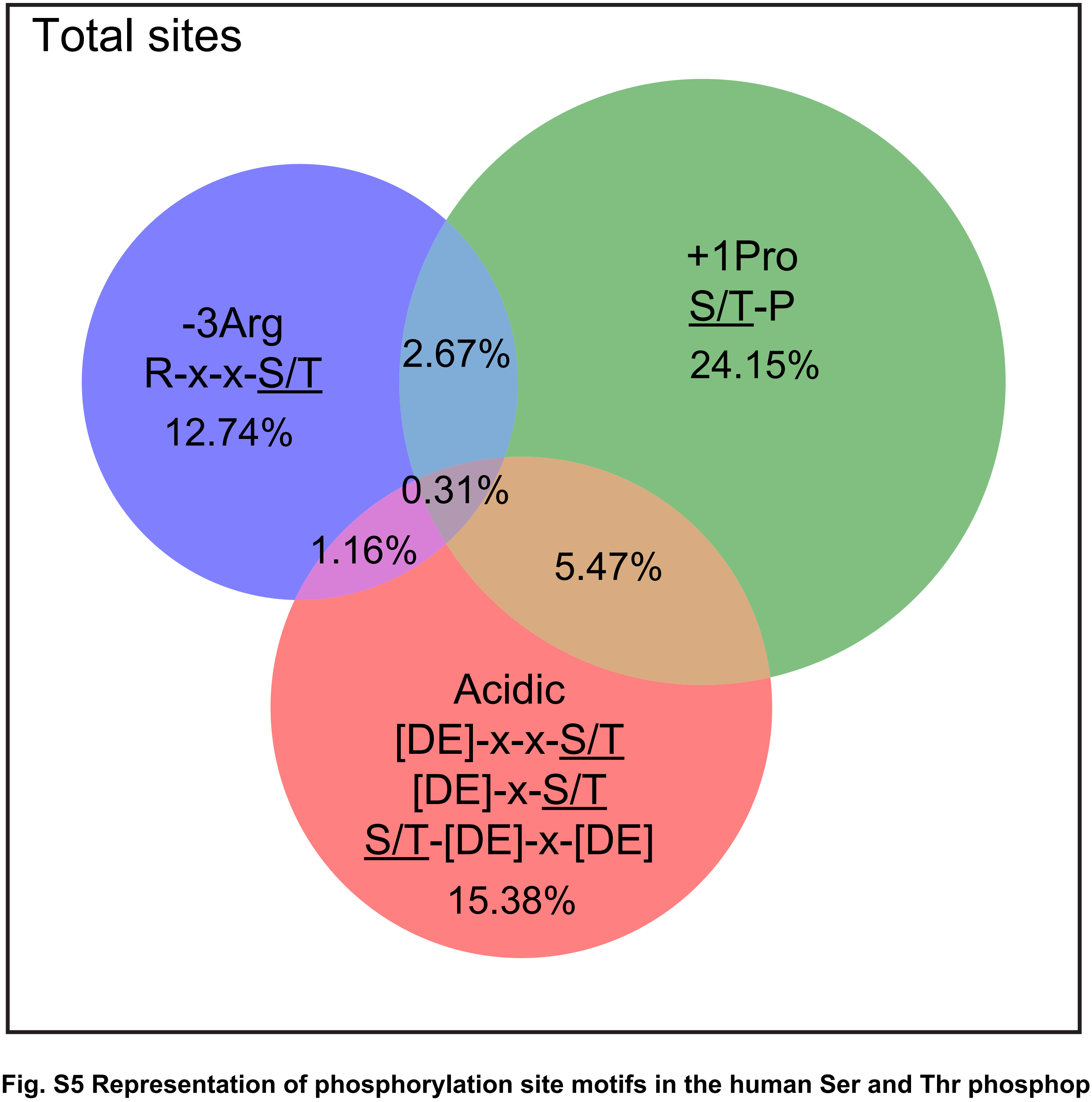


**Fig. S5 Representation of phosphorylation site motifs in the human Ser and Thr phosphoproteome.**
Venn diagram representation of the percentages of three prominant Ser/Thr kinase motif features, pertaining to Clusters 1, 2, and 3 in Figure 2, across ~50,000 human serine and threonine phosphorylation sites that have been detected in at least 5 independent high-throughput (mass spectrometry) experiments or reported in at least one low-throughput experiment. The phosphorylated residues in the logos are represented as S/T.

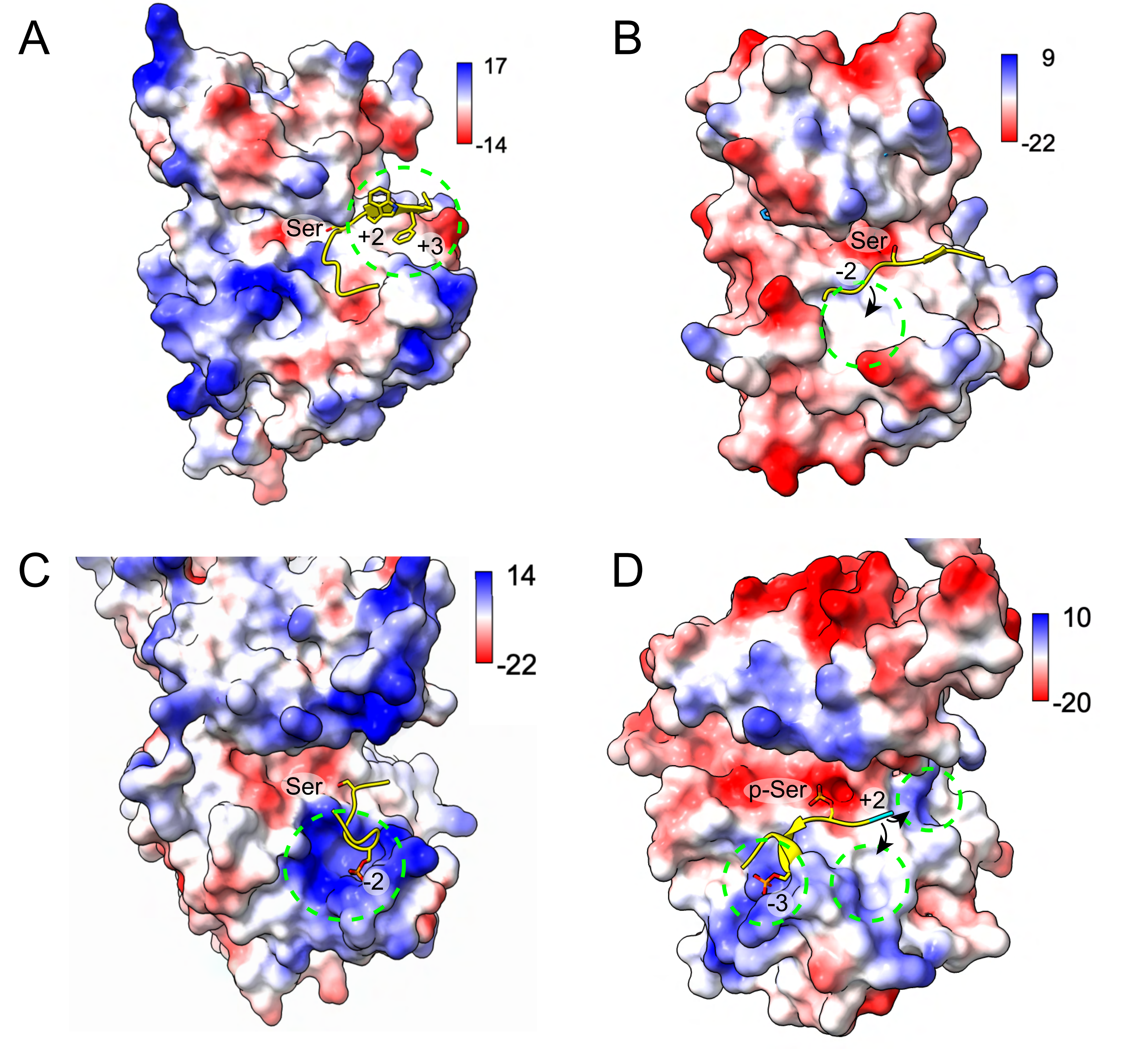


**Figure S6. Structural models of kinase-substrate complexes.**
(A) Synthetic peptide from its complex with PAK4 (PDB: 2Q0N) modeled onto WNK3 (PDB: 5O26). Dotted circle highlights a shallow hydrophobic pocket accommodating a +3 Phe residue. (B) GSK3 peptide from its complex with AKT2 (PDB: 1O6L) modeled onto CAMKK2 (PDB: 2ZV2). Circle indicates a hydrophobic pocket that could accommodate a -2 aliphatic residue. (C) Diphosphorylated peptide from p63 bound to CK1δ (PDB: 6RU6) modeled onto GRK2 (PDB: 1YM7). Circle shows positive surface potential in the vicinity of the -2 and -3 pSer residues. (D) The p63 peptide from (C) modeled onto YANK1 (PDB: 4FR4) showing potential binding sites for -3 and +2 phosphorylated residues. Surface electrostatics are represented with Coulombic potentialvalues were computed in ChimeraX and represented by scale bars (kcal/mol·e).

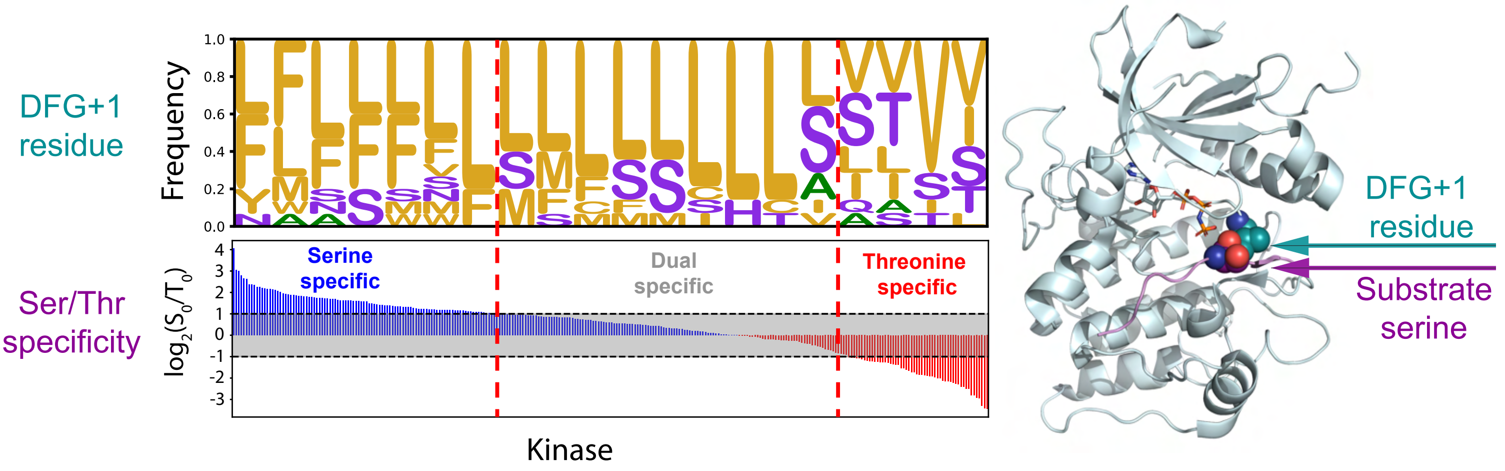


**Fig. S7 Gobal analysis of the relationship between the DFG+1 amino acid and preference for the serine versus threonine phosphoacceptor.**
(A) (bottom) Bars indicate relative preference for a Ser or Thr phosphoacceptor residue for each kinase, arranged in order of decreasing Ser/Thr selectivity. The frequency of amino acids at the DGF+1 positions of corresponding kinases is displayed above (bin size: 15 kinases). (B) Close proximity between DFG+1 residue and substrate phosphoacceptor residue shown on the structure of AKT1 bound to substrate peptide (PDB: 1O6K).

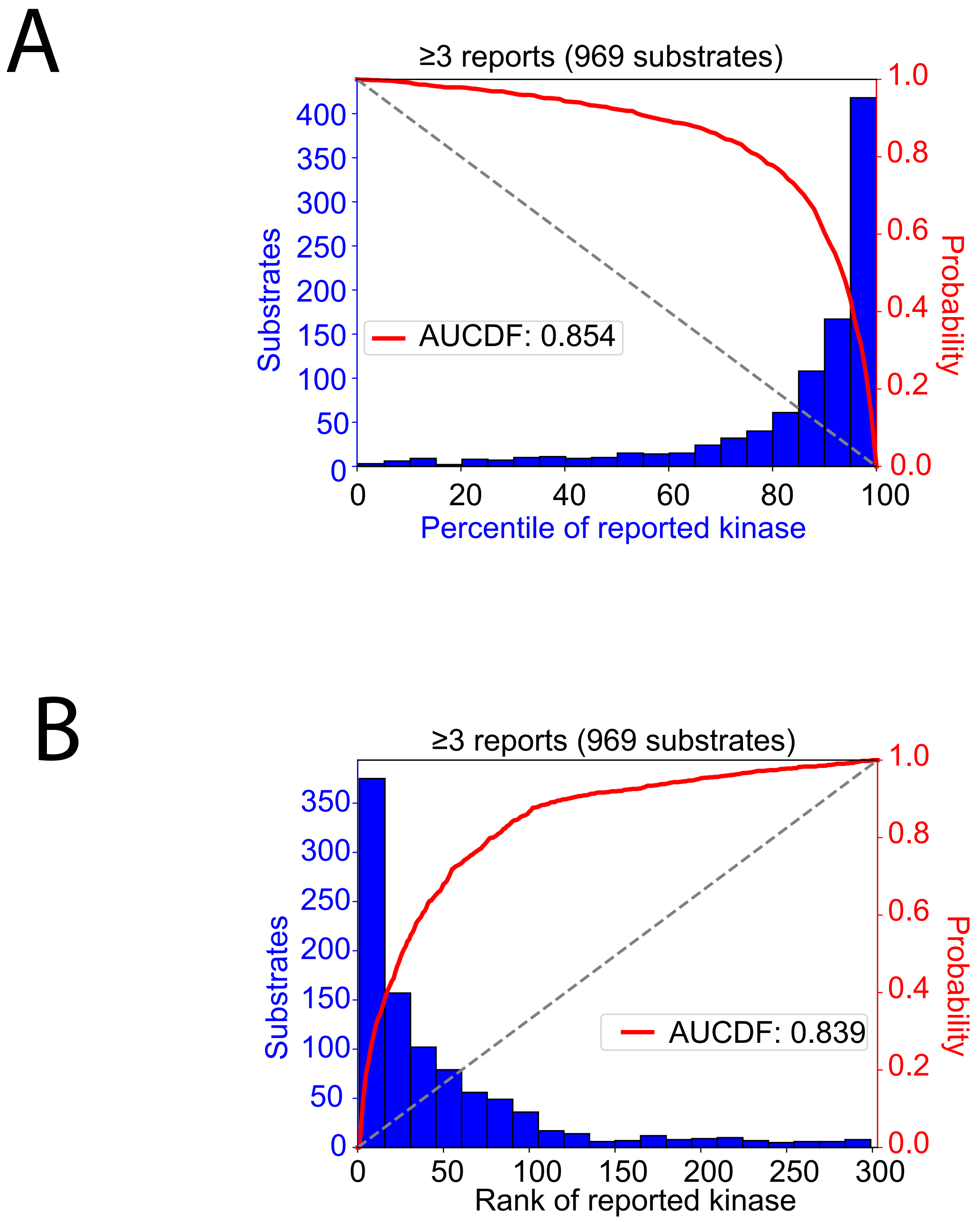


**Fig. S8 Global performance analysis of motif-based predictions.**
(A) Percentile distribution of substrates for their literature-annotated kinases. (B) Rank distribution of kinases for their literature-annotated substrates

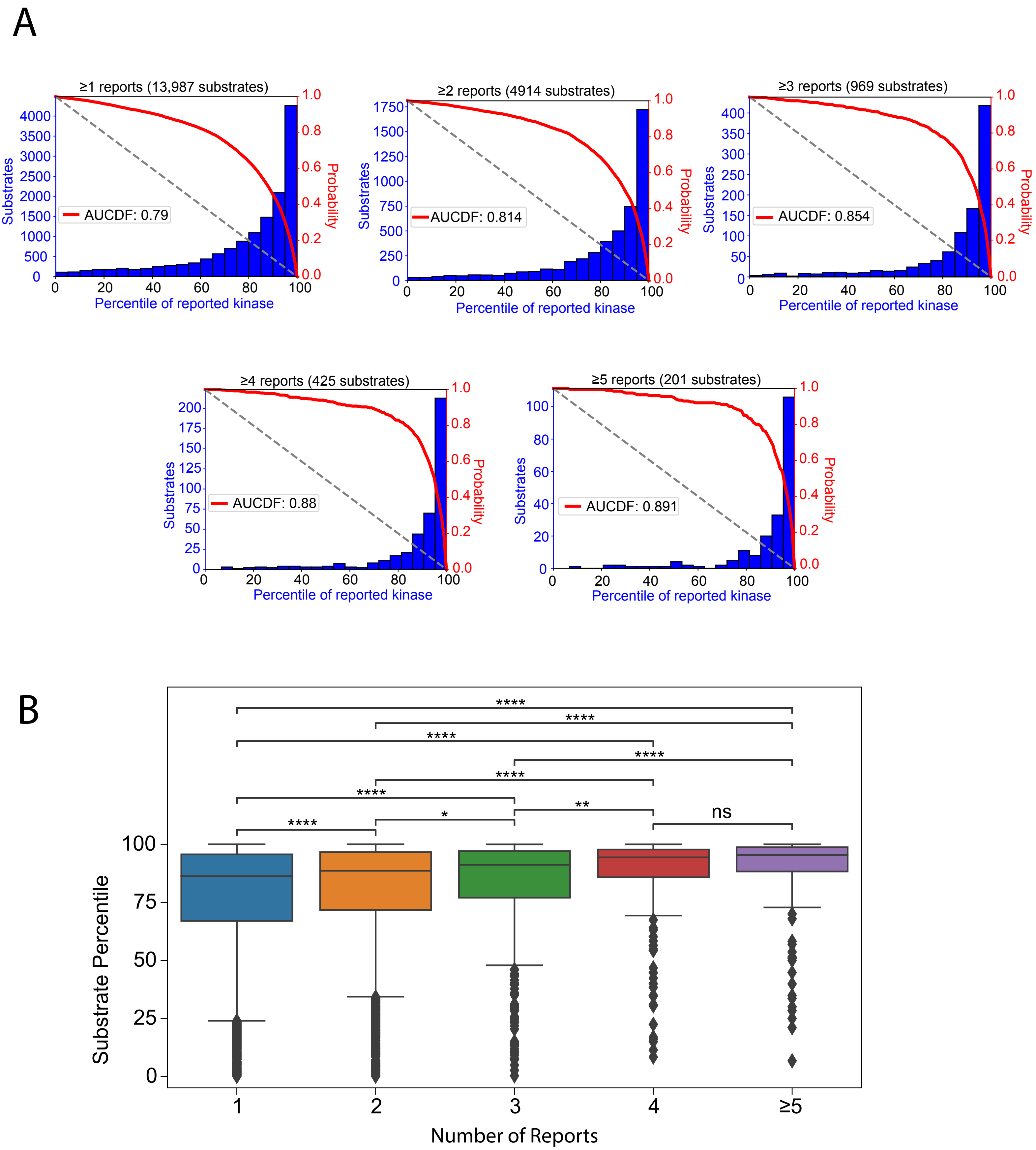


**Fig. S9 Global performance analysis of substrate percentile scores for their literature-annotated kinases.**

(A) Histogram of number of substrates as a function of the percentile-score of their literature-annotated upstream kinase. (B) Percentile-score of the literature-annotated kinase-substrate pairs, as a function of number of reports. The more reports the kinase-substrate pairs have, the higher the percentile-score of the reported kinase for its substrate.

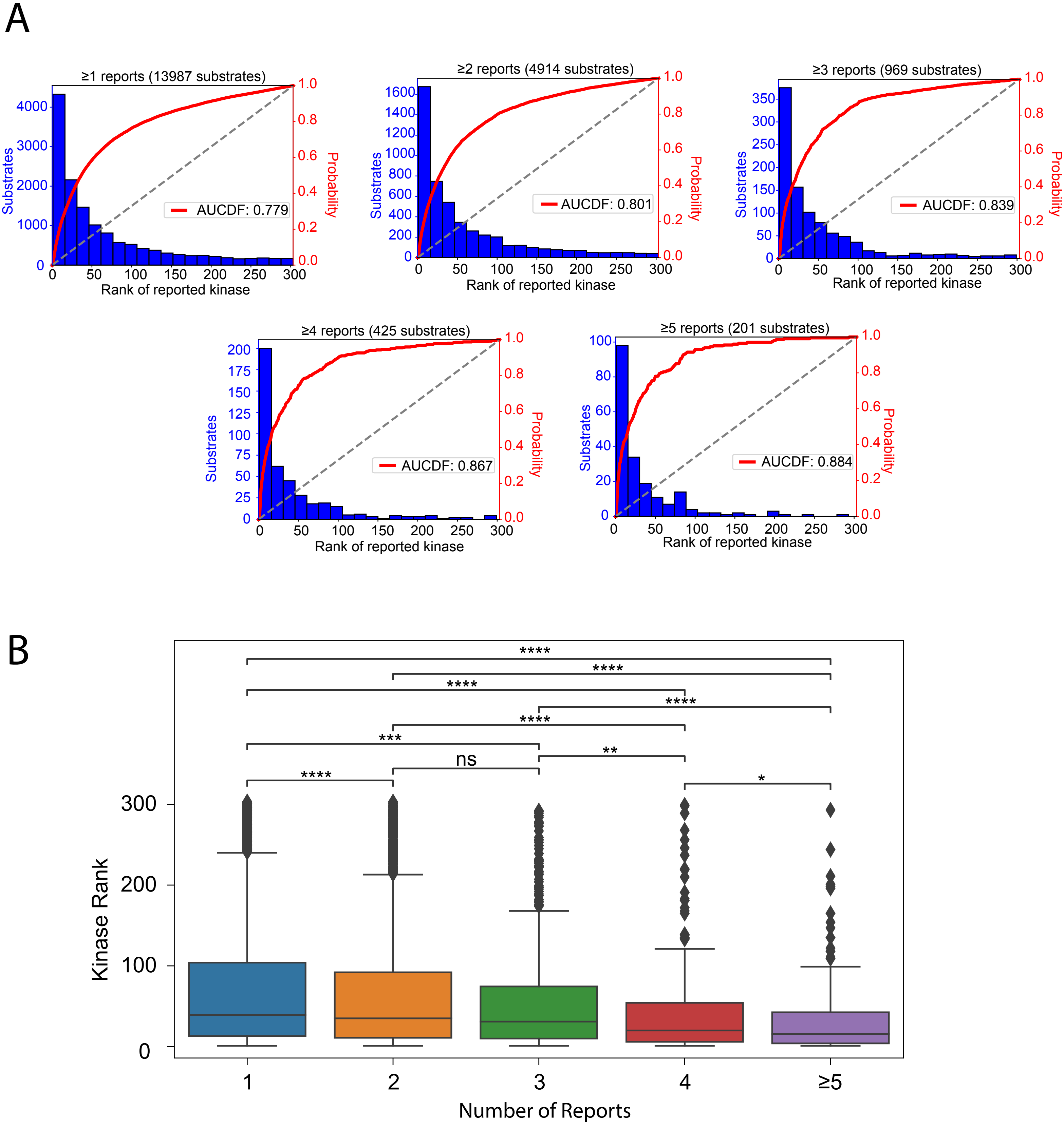


**Fig. S10 Global performance analysis of kinase ranks for their literature-annotated substrates.**

(A) Histogram of number of substrates as a function of the rank (based on percentile-score) of their literature-annotated upstream kinase. (B) Rank of the literature-annotated kinase-substrate pairs, as a function of number of reports. The more reports the kinase-substrate pairs have, the higher the reported kinase is ranked for its substrate.

A

### Inhibition of the pyruvate dehydrogenase complex by PDHK family kinases

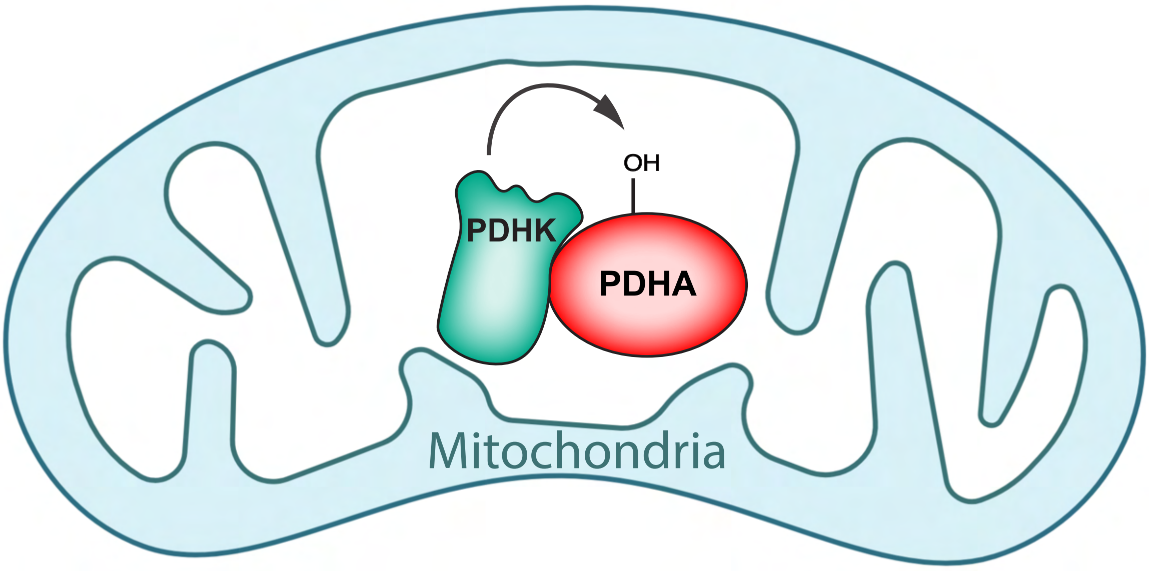


| Substrate: PDHA1 Ser293 | | |
|---|---|---|
| Phosphorylation site: RYHGH**S**MSDP | | |
| Rank | Kinase | Percentile |
| 1 | **BCKDK** | 99.99 |
| 2 | **PDHK1** | 99.82 |
| 3 | MASTL | 99.73 |
| 4 | **PDHK4** | 99.53 |
| 5 | TLK2 | 99.04 |
| 6 | GRK5 | 98.91 |
| 7 | HUNK | 98.56 |
| 8 | TTBK2 | 98.39 |
| 9 | TLK1 | 98.22 |
| 10 | TBK1 | 98.02 |

B

### Activation of ERK by MEK

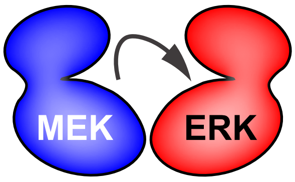


| Substrate: ERK1 Thr202/ERK2 Thr203 | | |
|---|---|---|
| Phosphorylation site: HTGFL**T**EYVA | | |
| Rank | Kinase | Percentile |
| 1 | **MEK2** | 99.73 |
| 2 | **MEK1** | 98.45 |
| 3 | YANK2 | 97.92 |
| 4 | MASTL | 97.91 |
| 5 | ASK1 | 96.84 |
| 6 | MEK5 | 96.74 |
| 7 | PRP4 | 95.54 |
| 8 | VRK2 | 94.97 |
| 9 | STLK3 | 94.66 |
| 10 | BMPR1A | 93.14 |

**Fig. S11 Motif-based scoring results for prominent kinase-substrate relationships.**
(A) Illustration of the mitochondrial-localized regulation of the pyruvate dehydrogenase complex through phosphorylation by the PDHKs. Scoring results for PDHA1 Ser293 highlighting PDHK family kinases. (B) Illustration of docking-driven activation of ERK1/2 through phosphorylation by the MEK1/2. Scoring results for sequence at ERK1 Thr202/ERK2 Thr203 highlighting MEK1 and MEK2.

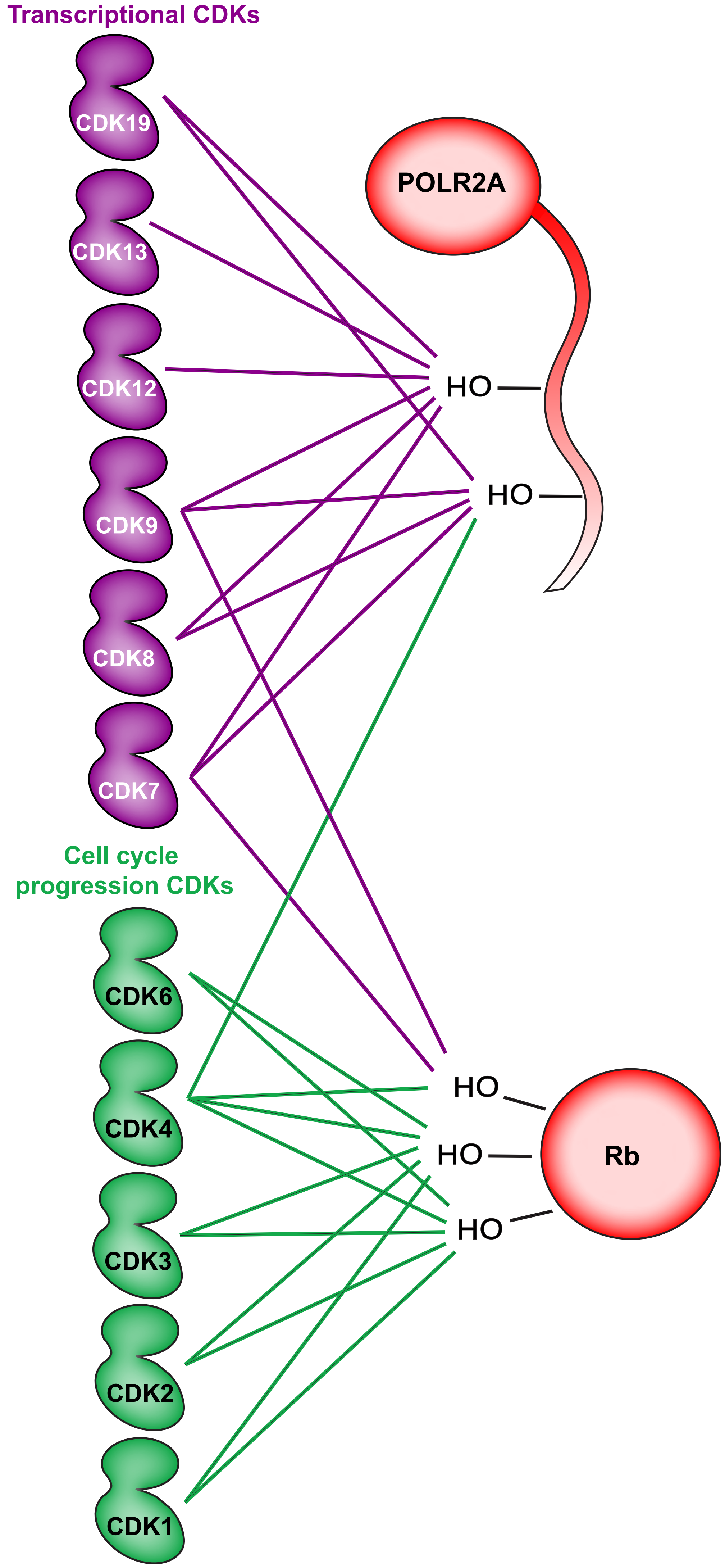


### POLR2A

| Substrate: POLR2A Ser1616 (ser2) | | |
|---|---|---|
| Phosphorylation site: QSPSYSPTSP | | |
| Rank | Kinase | Percentile |
| 1 | CDK7 | 94.69 |
| 2 | ERK5 | 89.64 |
| 3 | CDK13 | 89.47 |
| 4 | CDK12 | 88.35 |
| 5 | CDK19 | 88.33 |
| 6 | CDK8 | 86.78 |
| 7 | CDK9 | 86.48 |
| 8 | CDK17 | 85.58 |
| 9 | JNK3 | 85.32 |
| 10 | ERK2 | 85.17 |

| Substrate: POLR2A Ser1619 (ser5) | | |
|---|---|---|
| Phosphorylation site: SYSPTSPSYS | | |
| Rank | Kinase | Percentile |
| 1 | CDK7 | 99.15 |
| 2 | CDK8 | 98.36 |
| 3 | ERK2 | 98.15 |
| 4 | CDK19 | 98.07 |
| 5 | PDHK4 | 97.13 |
| 6 | ERK1 | 96.86 |
| 7 | CDK9 | 95.31 |
| 8 | P38A | 94.88 |
| 9 | CDK4 | 94.26 |
| 10 | P38B | 94.10 |

## Rb

| Substrate: Rb Ser780 | | |
|---|---|---|
| Phosphorylation site: RPPTLSPIPH | | |
| Rank | Kinase | Percentile |
| 1 | ERK5 | 98.54 |
| 2 | DYRK4 | 97.97 |
| 3 | CDK7 | 97.73 |
| 4 | CDK4 | 97.09 |
| 5 | DYRK1B | 96.94 |
| 6 | PRP4 | 96.76 |
| 7 | DYRK2 | 96.54 |
| 8 | CDK9 | 96.31 |
| 9 | BUB1 | 96.00 |
| 10 | KIS | 95.86 |

| Substrate: Rb Ser807 | | |
|---|---|---|
| Phosphorylation site: GNIYISPLKS | | |
| Rank | Kinase | Percentile |
| 1 | CDK2 | 97.05 |
| 2 | CDK6 | 94.81 |
| 3 | CDK3 | 93.06 |
| 4 | CDK4 | 89.65 |
| 5 | CDK1 | 87.13 |
| 6 | CDK5 | 82.58 |
| 7 | PINK1 | 80.20 |
| 8 | NLK | 76.13 |
| 9 | PRP4 | 74.46 |
| 10 | ERK2 | 73.74 |

| Substrate: Rb Ser811 | | |
|---|---|---|
| Phosphorylation site: ISPLKSPYKI | | |
| Rank | Kinase | Percentile |
| 1 | NLK | 99.17 |
| 2 | CDK1 | 98.41 |
| 3 | CDK3 | 98.36 |
| 4 | CDK2 | 98.31 |
| 5 | ERK2 | 97.30 |
| 6 | CDK4 | 97.10 |
| 7 | CDK5 | 96.94 |
| 8 | CHAK2 | 95.96 |
| 9 | ERK5 | 95.68 |
| 10 | CDK6 | 95.17 |

**Fig. S12 Delineation of the CDK subgroups by their motif scores for their specialized substrates.**
(Left) Illustration of site-specific phosphorylation of Retinoblastoma protein (Rb) and RNA Polymerse II (POLR2A) by their respective CDKs, cell cycle progression CDKs (green) and transcriptional CDKs (purple). Links between kinase and substrate correspond to favorable scoring results (shown on right).
